# Ferret model of bleomycin-induced lung injury shares features of human idiopathic pulmonary fibrosis

**DOI:** 10.1101/2025.05.08.652970

**Authors:** Shuang Wu, Ian Drive, Meihui Luo, Hikaru Miyazaki, Smitha Shambhu, Dimitry Popov, Liyuan Yang, Jing Wang, Jia Ma, Junfeng Guo, Jarron Atha, Aleksandra Tata, Eric A. Hoffman, Yujiong Wang, Purushothama Rao Tata, Martin B Jensen, John F Engelhardt, Vishwaraj Sontake, Xiaoming Liu

## Abstract

Idiopathic pulmonary fibrosis (IPF) is a debilitating lung disease with limited therapeutic options. The development of effective therapies has been hindered by the lack of models that recapitulate key features of human disease. Here we report a bleomycin-induced ferret PF model characterized by an irreversible decrease in pulmonary compliance and an increase of opacification, accompanied by “honeycomb cyst-like” structures and “proximalization” of distal lung epithelium. Cellular and molecular analysis by single-nucleus RNA sequencing revealed a significant shift in distal lung epithelium towards proximal epithelial phenotype. Importantly, a histopathological pattern of bronchiolization encompassing divergent atypical epithelial cells and KRT17^+^/TP63^+^/KRT5^low^ “basaloid-like” cells was present in the distal fibrotic lung lesions. Trajectory analysis revealed AT2 cells transition through multiple cell-states in bleomycin injured ferret lungs, particularly AT2 to KRT8^high^/KRT7^low^/SOX4^+^ and eventually to KRT8^high^/KRT7^high^/SFN^+^/TP63^+^/KRT5^low^ “basaloid-like” cells. Further, immunofluorescence analyses demonstrated KRT7 and KRT8 populations reside overlaying the ACTA2-positive myofibroblasts in fibrotic foci, implying their pro-fibrogenic activity similar to human IPF lungs. Collectively, our results provide evidence that bleomycin-induced lung injury in ferrets recapitulates pathophysiological, cellular, and molecular features of human IPF, suggesting that they may be a reliable model for understanding mechanisms of IPF pathogenesis and for testing therapeutic strategies for treatment of IPF.

**Take Home Message:** Bleomycin-induced acute lung injury in the ferret recapitulates pathophysiological, cellular, and molecular features of human IPF; thus the ferret may be a reliable species for studying mechanisms of IPF pathogenesis and testing therapeutic strategies.

## INTRODUCTION

Idiopathic pulmonary fibrosis (IPF) is a progressive pulmonary disease characterized by fibrotic lesions, with excessive deposition of extracellular matrix (ECM) and destruction of the lung architecture, which ultimately leads to decline of pulmonary function and respiratory failure with limited therapeutic options [1, 2]. The lack of clinically relevant and highly predictive models has greatly impeded our understanding of IPF pathogenesis, and consequently, the development of new treatments for IPF [3].

Although bleomycin-induced fibrosis in rodents share certain features of human IPF, most have failed to recapitulate the complex pathobiology, in particular the bronchiolization of distal alveoli and formation of honeycomb cyst-like structures that chronically persist. In general, lung fibrosis in rodents exhibits predominantly reversible features with self-limiting disease [4, 5]. However, recent studies indicate that very old mice may exhibit progressive and chronic fibrosis [6]. Bleomycin-injured large animals such as canines [7], sheep [8, 9], pig [10], and non-human primates (NHPs) [11] tend to recapitulate most manifestations of human IPF, however, the need for biosafety containment, widespread inaccessibility of transgenic lines, and costs of these models can be challenging.

Domestic ferrets (*Mustela putorius furo*) have been recognized as excellent models for medical research on a variety of respiratory diseases owing to similarities of airway anatomy, lung cell biology, and physiology between ferrets and humans [12]. These diseases include influenza and SARS-CoV-2 infections [13, 14], smoke-induced chronic obstructive pulmonary disease (COPD) [15], obliterative bronchiolitis (OB) [16-18], emphysema caused by alpha-1 antitrypsin deficiency [19], cystic fibrosis (CF) [20-22], and acute lung injury followed by pulmonary fibrosis (PF) [23-25]. Each of these ferret models demonstrates potential to inform disease mechanisms that are similar to those in humans. Recent advances in ferret genetic engineering have made it possible to create sophisticated conditional transgenic ferrets with multiple transgenes, allowing for both fate mapping and gene manipulation [26, 27]. These capabilities position ferrets as a valuable alternative species for investigating complex mechanistic questions in disease models.

Here we report a comprehensive evaluation of the histopathology, pulmonary function, and radiographic phenotypes, as well as cellular and molecular features, of a bleomycin-induced ferret model of PF. Our findings reveal that this bleomycin-induced ferret model exhibits key features of human IPF, suggesting it could serve as a reliable model for investigating the underlying mechanisms of IPF and accelerating the development of effective therapies.

## RESULTS

### Bleomycin induces histopathologic and biochemical features resembling human IPF in ferret lungs

Following three doses of bleomycin injury, a gradual loss of body weight was observed in both male and female ferrets with most animals reaching a humane endpoint before 12 weeks post bleomycin challenge (Fig. 1A-C). By 4-weeks post bleomycin challenge, lung histopathology revealed a patchy distribution of fibrotic foci and honeycomb cyst-like structures (Fig. 1D) with significant collagen deposition (Fig. 1E,F). Consistent with our histomorphometric findings, the hydroxyproline (HYP) content was significantly higher in the plasma and lungs of bleomycin ferrets after the challenge than in age- and sex-matched controls (Suppl. Fig. S1A-C). Plasma HYP content was also significantly higher in older bleomycin-injured ferrets than that in their younger counterparts (Suppl. Fig. S1D), while no differences were detected between males and females (Suppl. Fig. S1E). In agreement with the histopathological changes and increased matrix metalloproteinase-2 (MMP-2) and collagen content (Suppl. Fig. S1F,G), the Ashcroft scores were significantly higher in bleomycin ferrets compared to controls (Fig. 1G,J). No significant decrease in the Ashcroft scores were observed beyond the 4-week timepoint, but the loss of some ferrets to clinical endpoints led to a downward trend due to drop-out of the most severely diseased animals (Fig. 1G). Notably, older ferrets (≥11 months old) were more sensitive to bleomycin injury (i.e., had higher Ashcroft scores) than younger ferrets (≤10 months) (p = 0.0030, N = 7), but there were no differences between male and female ferrets (Fig. 1H,I). This age-related observation regarding PF is consistent with the incidence and prevalence of IPF in human populations [28, 29] and bleomycin-based models of IPF in other species [4, 6, 30].

**FIGURE 1.**
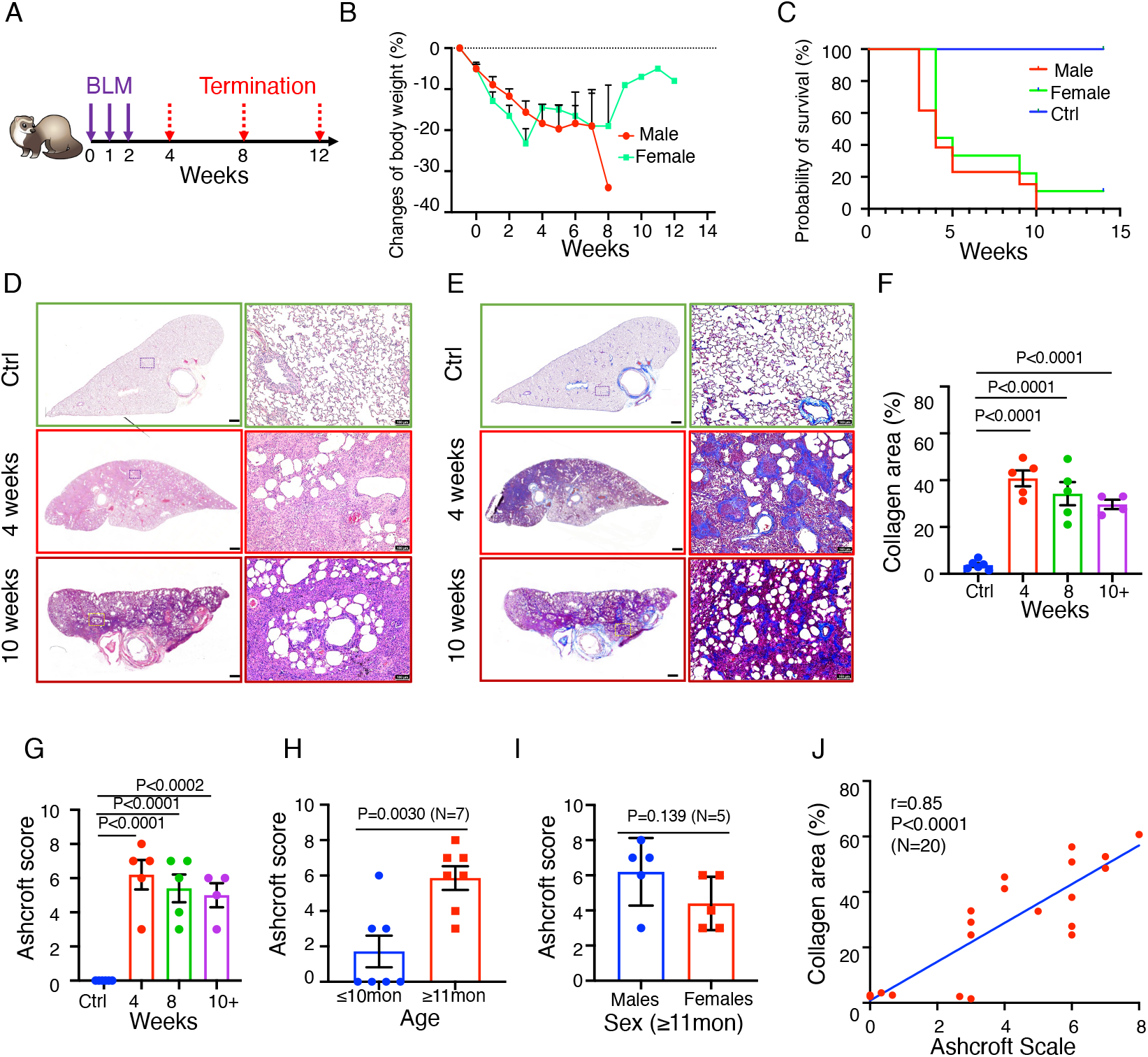
Bleomycin challenge in ferrets induces pulmonary fibrosis. (A) Schema representing the experimental timelines for bleomycin-induced PF in ferrets. Ferrets received 3 bleomycin (BLM) doses intratracheally at 1-week intervals (purple arrows), and lung tissues were harvested and analyzed at 4, 8, or 12 weeks (red arrows) after the first bleomycin challenge. Healthy age- and sex-matched ferrets treated with saline served as controls. (B) Curves of changes in body weight for male and female ferrets up to 13 weeks after the first bleomycin exposure. Data represents the mean ± SD. (C) Kaplan-Meier survival curve for male and female ferrets up to 14 weeks after the first bleomycin exposure. (D) Representative images of H&E-stained lungs from control and bleomycin-challenged ferrets at the indicated timepoints. (E) Representative images of Masson’s trichrome stained lungs (for collagen deposition) from control and bleomycin-challenged ferrets at the indicated timepoints. Images in right panels of D and E are enlargements of boxed areas in the images at left. (F) Quantification of % collagen area as determined from Masson’s trichrome-stained images from control and bleomycin-injured ferrets at the indicated timepoints. (G) Ashcroft scores in bleomycin-challenged ferrets at 4, 8 and 10+ weeks. (H) Ashcroft scores in ferrets ≤10 months and ≥11 months old, following challenge with bleomycin (P=0.0030, N=7). (I) Ashcroft scores in male vs. female ferrets challenged with bleomycin (P=0.139, N=5). (J) Collagen deposition as a function of Ashcroft score in lungs from ferrets challenged with bleomycin, as determined by analysis of the Pearson correlation coefficient (r) (r=0.85, P<0.0001, N=20). Data in F-I represent the mean ± SEM. Statistical significance was calculated using One-way ANOVA, followed by Dunnett’s comparison test, with (N) denoted in the graph. Data for individual ferrets are represented by dots. Scale bars in D and E equal 2 mm (left panels) or 100μm (right panels).

In addition to the histopathological similarity between the bleomycin-challenged ferret and human IPF phenotype, biochemical and molecular analyses revealed that the profile of circulatory biomarkers in bleomycin ferrets is similar to that in people with IPF. Specifically, we found the protein levels for known human IPF biomarkers (KL-6, SFTPA, SFTPD, MUC5AC, MUC5B, MMP-1, MMP-2, MMP-3, and MMP-7) were significantly higher in the plasma of bleomycin-challenged ferrets compared with saline controls (Suppl. Fig. S2); the one exception was S100A12. Of note, the highly-studied epithelial-specific circulatory biomarker KL-6, which is used to distinguish between IPF and other common forms of interstitial lung disease (ILD) [31, 32], was more abundant in the plasma of older bleomycin ferrets than their younger counterparts without gender differences (Suppl. Fig. S3A,B). The expression of SFTPA was also slightly higher in bleomycin ferret lungs related to control, as determined by immunostaining and immunoblotting assays (Suppl. Fig. S3C-E).

### Radiographic and pulmonary function assays reveal clinical manifestations of fibrosis in bleomycin-injured ferret lungs

Consistent with the histopathological findings, CT lung imaging showed diffuse homogeneous areas of increased lung opacification. The high attenuation area (HAA) was greater in the right vs. left lung (Suppl. Fig. S4A-C), likely due to uneven distribution of bleomycin during intratracheal delivery (Fig. 2A-C, Suppl. Fig. S4D,E). Consistently, pulmonary function testing (PFT) showed that the inspiratory capacity (IC) was significantly lower in both male and female ferrets at 4 weeks post-bleomycin challenge as compared to baseline (Fig. 2D; Suppl. Fig. S5A,B). Using FEV_0.4_ as a ferret equivalent to human FEV_1_ [19], we observed an restrictive pulmonary phenotype in males that was significant with an increased FEV_0.4_/FVC (Fig. 2E). Other PFT metrics, including quasi-static compliance (ml/cmH_2_O) (Cst) (Suppl. Fig. S5C) and dynamic compliance of the respiratory system (Crs, ml/cmH2O) (Suppl. Fig. S5D), were significantly decreased in ferrets at 4 weeks post bleomycin challenge. Elastance of the respiratory system (Ers, cmH2O/ml) (Suppl. Fig. S5E) and resistance (Rrs) (Suppl. Fig. S5F) were moderately increased. The decrease in Cst confirms that lung compliance in bleomycin ferret lungs is reduced (Suppl. Fig. S5C). Consistent with the more advanced fibrosis observed in male bleomycin ferrets, these animals demonstrated a significantly greater reduction in IC than their female counterparts (Fig. 2F), whereas the increase of FEV_0.4_/FVC ratio was not significantly different between sexes (Fig. 2G). In addition, there was no significant difference in IC (Suppl. Fig. S5G) and FEV_0.4_/FVC ratio (Suppl. Fig. 5H) between younger (≤10 months) and older (≥11 months) ferrets (Suppl. Fig. S5G). Notably, the decline in lung function was strongly associated with the degree of pathohistological change in the lungs of bleomycin-induced ferrets, as suggested by the correlation between the change in IC (ΔIC) and PF as measured by the Ashcroft scores (r= -0.8486; 95% CI, -0.9485 to -0,5950, p<0.0001, n=15) (Fig. 2H). Also, the decrease of IC (Fig. 2I) and increase of FEV_0.4_/FVC ratio (Fig. 2J) did not recover to baseline levels by 12 weeks, indicating a sustained decline in respiratory function in these ferrets.

**FIGURE 2.**
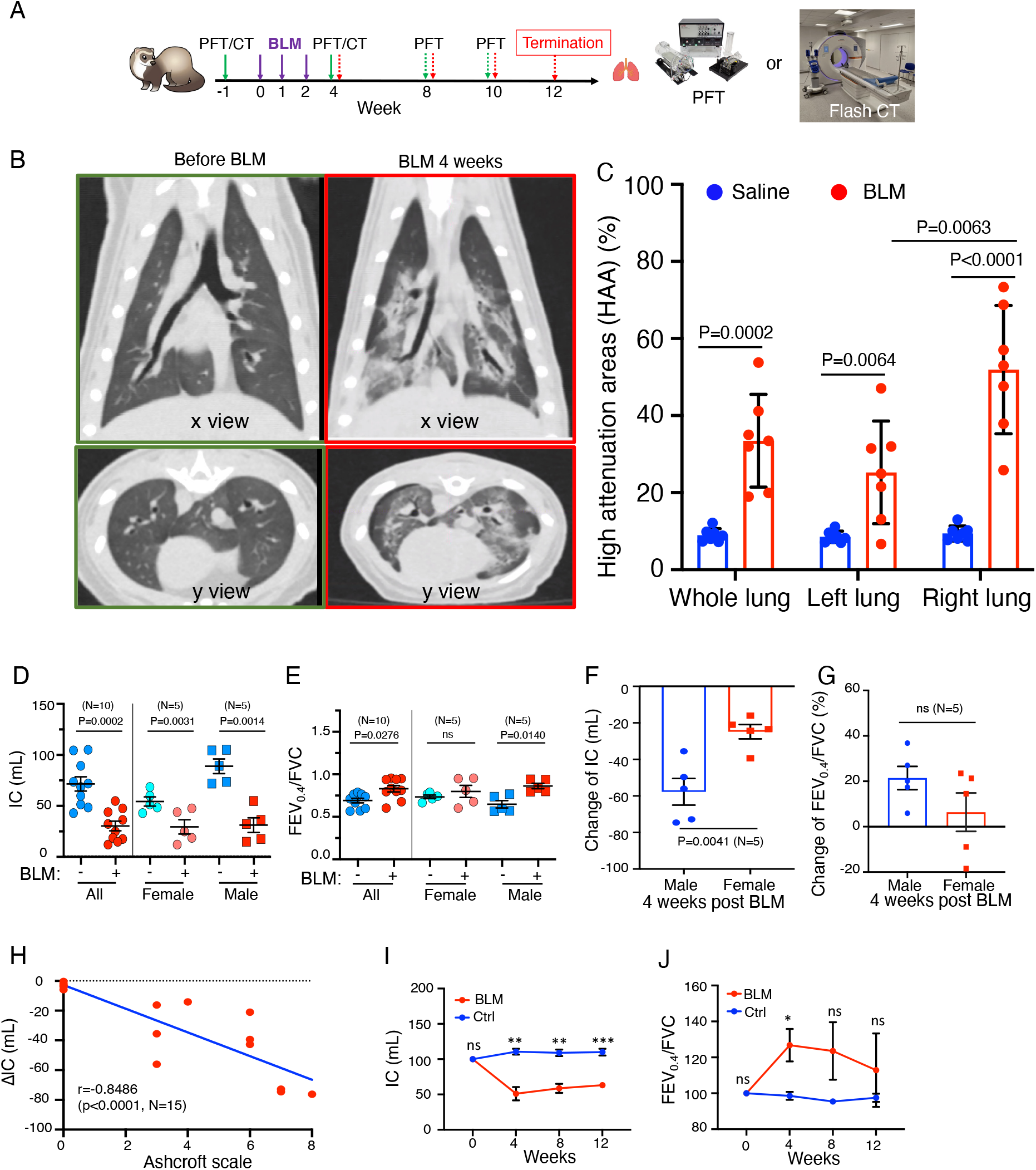
Bleomycin challenge in ferrets leads to decreased pulmonary compliance and restrictive pulmonary physiology. (A) Schema showing the timing of pulmonary function test (PFT) and computerized tomography (CT) in ferrets. PFT or CT was performed at one week (−1 week) before and at the 4, 8, and/or 10 weeks timepoints. (B) Representative 2D images (x view, upper panels; y view, lower panels) of a ferret at the indicated timepoints. (C) Quantification of radiographic opacity in lung, as determined by the percentage of high-attenuation areas (HAA) in whole lung and in the separated left and right lungs. (D,E) Bleomycin-induced changes in inspiratory capacity (IC, mL at 30 cmH_2_O) (D) and FEV_0.4_/FVC ratio (E) at 4 weeks in 10 animals (5 males and 5 females) in which PFT was examined at one week before and 4 weeks after the bleomycin challenge. (F,G) Differences in IC (F) and FEV_0.4_/FVC ratio (G) by sex. (H) Correlation between IC (*Δ*IC) and Ashcroft score as determined by analysis of Pearson correlation coefficient (r) (r=-0.8486, P<0.0001, N=15). (I,J) Differences in IC and FEV_0.4_/FVC ratio in control vs. bleomycin-challenged ferrets analyzed through 12 weeks (control N=3, bleomycin N=4): Data were analyzed in sex-mixed group and sex-specific differences are indicated in figure labels for D-H. Unpaired *t* test with the number of samples (control N=3, bleomycin N=4) of samples denoted in the graph for I and J.

We next evaluated age-dependent impaired pulmonary function longitudinally in a subset of bleomycin-injured ferrets, with more moderate disease progression that did not reach the humane terminal endpoint prior to study completion. In four ferrets that were challenged with bleomycin at the ages of 3-11 months, the pressure-volume (PV) loop at the last experimental timepoint was shifted downward at 4 weeks post-bleomycin challenge, and it gradually shifted upward in three younger animals (3-8 months) at 6-9 weeks post-bleomycin challenge (Suppl. Fig. S5I-K). In the fourth ferret (11-months-old at the first bleomycin exposure), the PV loop was stable at 6- and 11-weeks post-bleomycin challenge, and this animal met the humane terminal endpoint prior to 12-weeks (Suppl. Fig. S5L). It is worth noting that in all four of the longitudinally monitored bleomycin-injured animals, the PV loop did not recover to the initial baseline level (Suppl. Fig. S5I-L). Decline in pulmonary function in these bleomycin-induced IPF ferrets was further corroborated by the histopathological changes in lungs (Suppl. Fig. S5I’-L’), and other PFT parameters, including IC, Rrs, FEV_0.4_/FVC, Crs, Ers, and Cst (Suppl. Fig. S5M). These data suggest that impaired pulmonary function is persistent in bleomycin-induced ferrets up to 12 weeks, but that younger animals may have a greater capacity to recover from injury.

### Emergence of diverse aberrant epithelial cell phenotypes reminiscent of human IPF in bleomycin-injured ferret lungs

To gain insight into cell-specific alterations in bleomycin-induced ferret lungs, we performed single nucleus RNA-sequencing (snRNA-Seq) analysis (Fig. 3A). We profiled 114,125 cells from the distal lungs of 9 ferrets. Among these, 66,864 cells were obtained from 5 bleomycin-injured ferrets with Ashcroft scores of 4 or above, and 47,261 were obtained from 4 saline controls (Fig. 3A). Following the removal of batch effects across samples, all major cell types were annotated using human lung cell atlas (HCLA) reference annotations: epithelial, mesenchymal, endothelial, and immune cells (Fig. 3B) [33-35]. We identified 33 discrete cell clusters based on the presence of distinct marker sets (Fig. 3C,D). The unique expression profiles of signature genes in each main cell type were used for supervised clustering (Fig. 3E,F and Suppl. Fig. S6).

**FIGURE 3.**
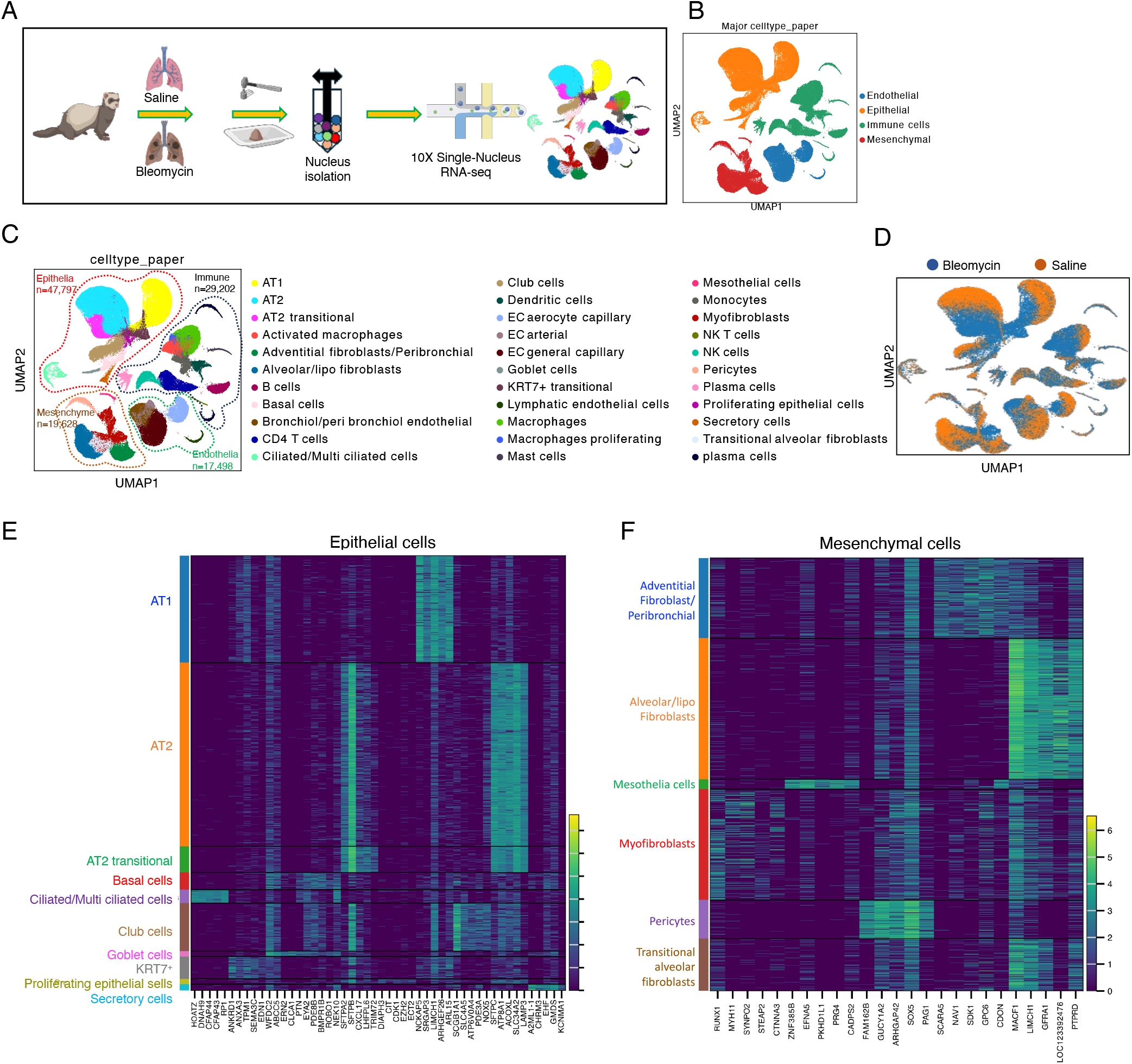
Single nucleus RNA sequencing (snRNA-Seq) reveals diverse aberrant epithelial and mesenchymal cell phenotypes in bleomycin-injured ferret lungs. (A)Schema showing workflow for generating and analyzing lung snRNA-Seq data. (B) UMAP of major cell classes (epithelial, endothelial, mesenchymal, and immune) captured in snRNA-seq analysis. (C) UMAP of all cells profiled; the 33 clustered cell types were annotated using human cell reference annotations from the HCLA. (D) UMAP showing the distribution of cell-types in bleomycin-injured (blue) and saline-treated (orange) ferret lungs. (E,F) Heatmap of marker genes used for classifying epithelial (E) and mesenchymal (F) cell types. Each cell type is represented by the top 5 genes ranked by Wilcoxon rank sum test for each cell type against the other cell types.

We next performed more focused analyses of epithelial cell changes in bleomycin-injured lungs and compared them to control counterparts. We identified 10 distinct cell types/states, encompassing typical epithelial cell types (e.g., AT1, AT2, basal, goblet, ciliated, and club), and epithelial cell states identified in human IPF lungs and rodent models (e.g., transitional AT2 cells), that express their unique marker genes in both bleomycin- and saline-treated ferret lungs (Fig. 4A,B,C). UMAP plots showed cell-type specific marker genes *SFTPC* (AT2), *AGER* (AT1), *MUC5B* (goblet cells in proximal airway and submucosal glands), *SCGB1A1* (club cells), and *SCGB3A2* (secretory cells) in corresponding epithelial subclusters (Fig. 4D).

**FIGURE 4.**
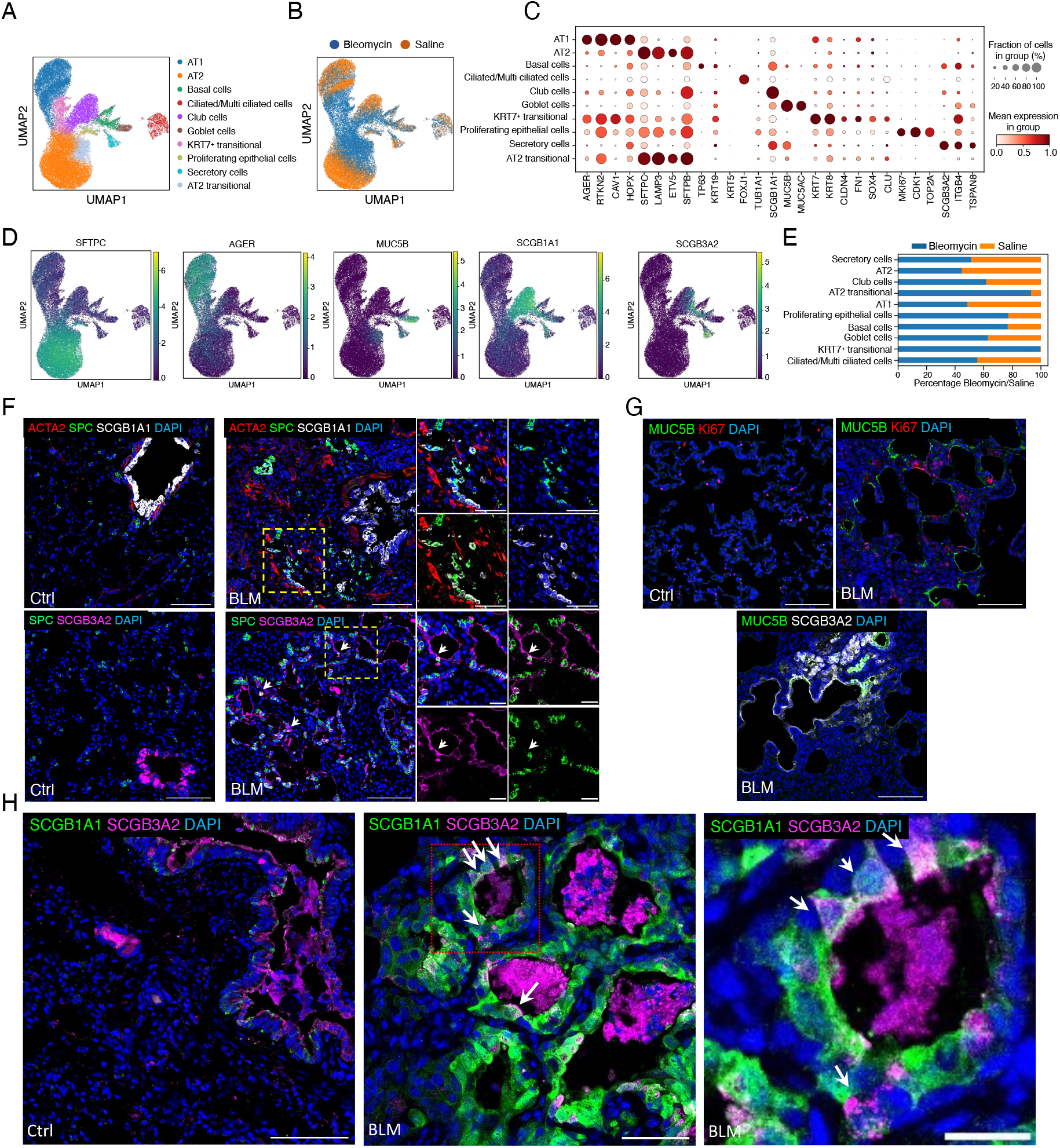
Bleomycin challenge leads to aberrant epithelial cells. (A) UMAP of snRNA-Seq visualizing 10 subclusters of epithelial cells in both BLM- and saline-treated ferret lungs.(B) UMAP of snRNA-Seq visualizing differences between epithelial cell-types in bleomycin- and saline-treated ferret lungs. (C) Dot plot showing frequency of markers of cell types and transitional cell states. (D) UMAPs visualizing the distribution of expression of the *SFTPC, AGER, MUC5B, SCGB1A1,* and *SCGB3A2* genes. (E) Bar plots showing the proportion of each epithelial-cell subtype enriched in bleomycin- and saline-treated lungs. (F) (Top panels) Representative immunofluorescence (IF) images showing cells that co-express the SPC (green) and SCGB1A1 (white) proteins and their proximity to ACTA2 (red)-expressing myofibroblasts. (Bottom panels) expression of SPC (green) and SCGB3A2 (megenta) (bottom panels) proteins in fibrotic lesions. Insets indicate enlarged region and individual channels for regions marked by yellow dashed line box. DAPI-nuclei (blue). (G) Representative IF images of MUC5B- and Ki67-stained (Top panels). Representative image of bleomycin-induced fibrotic regions co-staining for MUC5B and SCGB3A2 (Botton panel). (H) Representative IF images of saline- and bleomycin-treated tissue showing cells co-expressing SCGB3A2 (magenta) and SCGB1A1 (green). Insets indicate enlarged region and individual channels of regions marked by yellow dashed box. DAPI-nuclei (blue). Scale bars equal 100µm (main images), 50 µm in enlarged image in (F), and 20 µm in enlarged images in (H).

Analysis of differences in relative proportion of epithelial cell composition revealed a decrease in the number of AT1 (51.41% in saline vs. 48.59% in bleomycin) and AT2 (55.43% in saline vs. 44.57% in bleomycin) cells, and dramatic increases in the number of transitional AT2 cells (6.8% in saline vs. 93.2% in bleomycin) and KRT7^+^ cells (0.49% in saline vs. 99.51% in bleomycin) (Fig. 4E). The relative proportions of club (38.3% in saline vs. 61.7% in bleomycin), basal (23.25% in saline vs. 76.75% in bleomycin), goblet (37.12% in saline vs. 62.88% in bleomycin), and ciliated (44.61% in saline vs. 55.39% in bleomycin) cells were higher in the bleomycin compared to control lungs, indicating the “proximalization” of distal lung epithelium in ferret lung fibrosis (Fig. 4E). Corroborating the snRNA-seq data, immunofluorescence staining (IF) showed that the numbers of SCGB1A1 and SCGB3A2 expressing cells were significantly increased in fibrotic lungs in ferrets (Fig. 4F). Notably, in bleomycin ferret lungs a subset of SCGB1A1^+^ and SCGB3A2^+^ cells also expressed the AT2 cell marker SFTPC (SPC) (Fig. 4F), indicating that these putative transitional AT2 cells were phenotypically similar to previously described terminal and respiratory bronchiole (TRB)-specific AT0 cells [11], or respiratory airway secretory (RAS) cells [36]. Further, we found SCGB1A1^+^SCGB3A2^+^ double positive cells in “bronchiolized” regions (Fig. 4H), which are phenotypically similar to human TRB secretory cells (TRB-SCs) in both human IPF and bleomycin-injured primate lungs [11]. Note that these cells did not separately cluster in our single-cell analysis, presumably because they are few in number and share many of markers with AT2 and airway club/secretory cells. Collectively, single-cell and in situ validation data suggest that the lung fibrosis phenotype in ferrets emulates the AT0/RAS and TRB-SC dynamics that are observed in human IPF but not in other fibrosis models.

Genetic studies have demonstrated that MUC5B variants are strongly associated with the development of IPF and are considered an important risk factor [37-39]. Consistent with other scRNA-Seq analyses of human lungs [33, 34, 40, 41], we observed a significant increase in the MUC5B^+^ cell population in fibrotic lungs in ferret compared to controls (Fig. 4G). Further, IF staining revealed the presence of these cells in “honeycomb-like cyst” (Fig. 4G) [24]. The MUC5B^+^ cells were not proliferative, as evidenced by negative staining for the marker Ki67. However, most of the MUC5B^+^ cells also expressed SCGB3A2 in distal lung regions (Fig. 4G), consistent with the findings that a MUC5B risk variant promotes muco-secretory cell differentiation in IPF [39].

### Bleomycin induces bronchiolization of distal lung parenchyma in ferret lungs

Bronchiolization is a hallmark of the human IPF lung. It is characterized by ectopic aberrant basaloid epithelial cells and honeycomb cystic structures in the airspace of distal lung [33, 35]. Our snRNA-seq analysis showed an increase in the proportion of basal cells and their markers in fibrotic ferret lungs (Fig. 4E,5A). IF analysis for basal cell markers KRT17, TP63 and KRT5 further revealed a subpopulation phenotypically similar to basaloid cells (KRT17^+^/TP63^+^/KRT5^low^) within “bronchiolization-like” structures in the periphery of bleomycin ferret lung but not in the control ferret lung (Fig. 5B-D,D’). The *KRT17* gene is not annotated in the ferret genome and thus the analogous cell could not be interrogated at the single cell level, however, we used KRT7 as a surrogate marker for the presence of aberrant “basaloid-like” cells in bronchiolized regions and within the single cell dataset. Notably, the expression of KRT7 was observed in AT1 and AT2 cells of the alveolar space of bleomycin ferret lung but not that of the control lung (Suppl. Fig. S7A,B). Similar to the KRT17^+^/TP63^+^/KRT5^low^ population, a subset of TP63^+^ cells were also KRT7^+^/KRT5^low^ and similarly observed in the periphery of the bleomycin-treated lungs (Fig. 5E-G,G’). Further, the KRT7^+^ cells overlapped with the expression of KRT8, and subset of KRT7^+^/KRT8^+^ cuboidal cells express TP63 (Fig. 5H-I, I’). Similar to the observed KRT7^+^/TP63^+^/KRT5^low^ cells, we also found rare KRT8^+^/TP63^+^/KRT5^low^ cells and KRT7^+^/KRT17^+^/TP63^+^ cells (Suppl. Fig. S8) in fibrotic regions of bleomycin-injured ferret lungs. Taken together, these results suggest that KRT17^+^/TP63^+^/KRT5^low^, KRT7^+^/TP63^+^/KRT5^low^, and KRT8^+^/TP63^+^/KRT5^low^ cells of bleomycin-injured ferret lungs share features of basaloid cells observed in human IPF [33, 34]. Thus, these cells may contribute to the “bronchiolization-like” pattern observed in bleomycin-treated ferret lungs.

**FIGURE 5.**
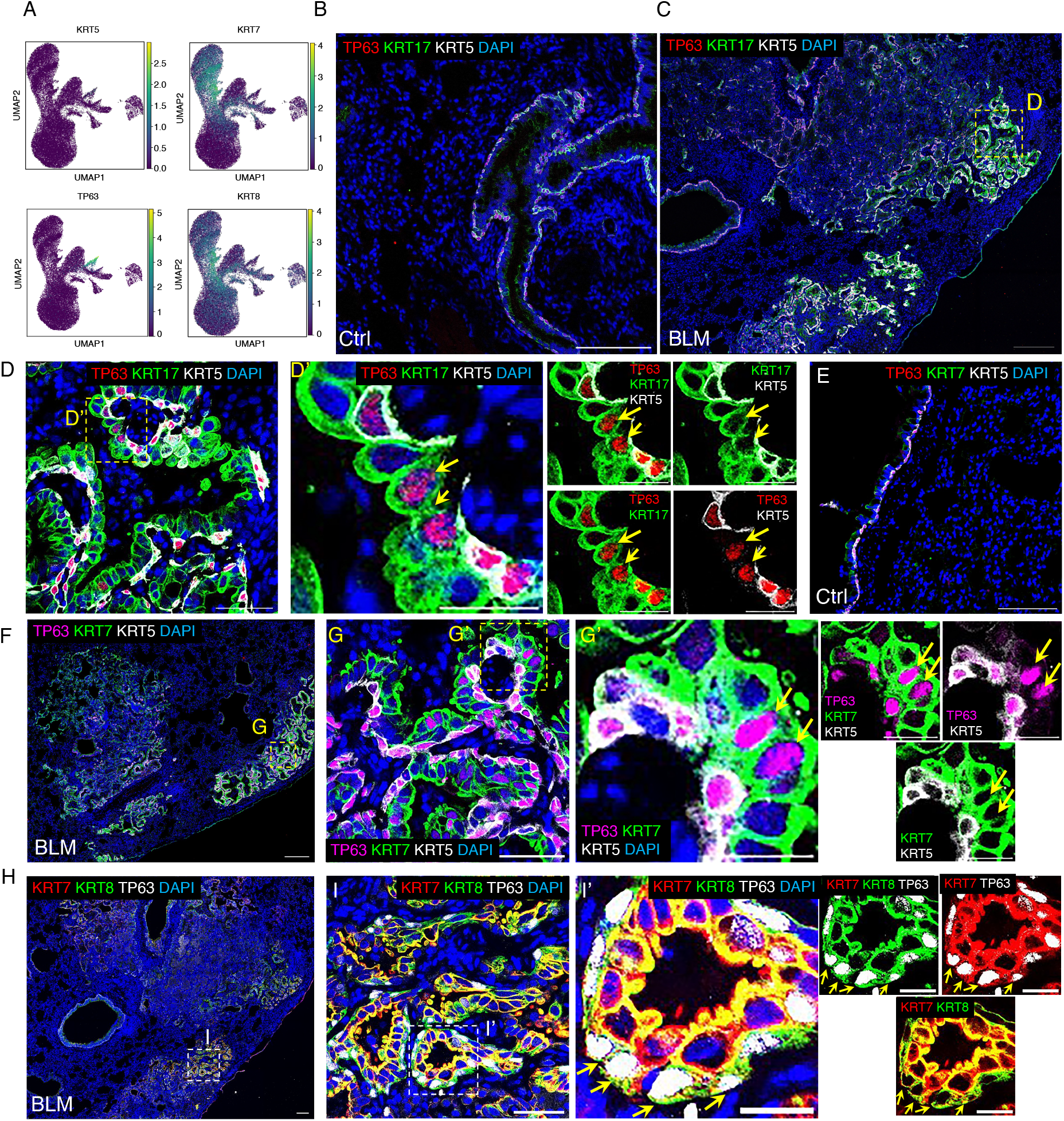
Bleomycin challenge leads to bronchiolization in the distal lung of ferret. (A) Heatmap showing the expression of epithelial-cell markers of proximal airways, *TP63, KRT7, KRT5 and KRT8* in ferret distal lungs. (B,C) Representative IF images of co-staining of KRT5 (white), KRT17 (green), and TP63 (red) in distal lung of control (B) and bleomycin (C) ferrets. (D) Inset from (C), showing bronchiolar morphology with KRT17 (green), KRT5 (white), and/or TP63 (red) -positive epithelial cells. (D’) Enlarged image of boxed area in panel D. Double-channel images in right panels of D’ showing cells that are positive for KRT17 and TP63 (red) but negative or low for KRT5 (white) (basaloid cells; arrows). (E,F) Representative IF images of co-staining of KRT5, KRT7, and TP63 in control (E) and bleomycin (F) ferret distal lungs. (G) Enlargement of boxed area in (F), showing bronchiolar morphology with KRT7, KRT5 and/or TP63-positive epithelial cells. (G’) Enlargement of boxed area in (G). Double-channel images in right panels of G’, showing cells that are positive for KRT7 and TP63 but negative or low for KRT5 (basaloid cells, arrows). (H-I) Representative IF images of co-staining of KRT7, KRT8, TP63 in distal lungs of bleomycin ferrets, showing aberrant KRT7 and KRT8 basaloid cells. (I) Enlargement of boxed area in (H), showing bronchiolar epithelium with KRT7, KRT8, and/or TP63-positive cells. (I’) Enlargement of boxed area in (I). Double channel images in right panels of I’ showing KRT7/KRT8/TP63 triple-positive cells (arrows). Scale bars in B, E equal 100 μm; in C,F,H equal 200 μm; in D,G,I are equal 50 μm; in D’,G’,I’ equal 20 μm.

To investigate the relationship between the KRT17- and KRT7-expressing basaloid-like cells (TP63^+^/KRT5^low^), we performed co-immunolocalization of KRT5, KRT7 and KRT17. As the progression of fibrotic lesions gradually progresses from the lung periphery toward the center of the lobe (Fig. 6A) [42, 43], we examined the expression of these keratins across these regions (Fig. 6B, B1-5). In the central regions of the bleomycin-treated lung, we observed simple squamous and cuboidal KRT7^+^ cells lacking KRT17 and KRT5 expression (Fig. 6B, B1). These KRT7^+^ cells gradually adopted KRT17 and/or KRT5 expression as lesions progressed to the peripheral lung, where the dysplastic epithelium retained a heterogeneous pattern of keratin expression within cuboidal and columnar cells (Fig. 6B, B1-B5). Of note, the co-expression of these three keratins overlapped mainly in columnar epithelial cells of lesions midway between the center and periphery (Fig. 6B4). The expression of KRT7 and KRT17 in columnar cells of the peripheral bleomycin lung consistently overlapped; however, a subset of KRT5^+^ basal-like cells did not express KRT7 or KRT17 (Fig. 6B5). Subpopulations of KRT7^+^/KRT17^+^ cells also expressed TP63, suggesting they had adopted a “basaloid-like” phenotype (Fig. S8D-F). In addition, in the bleomycin-treated ferret lung is highly heterogenous, as in human IPF [33, 34, 44, 45].

**FIGURE 6.**
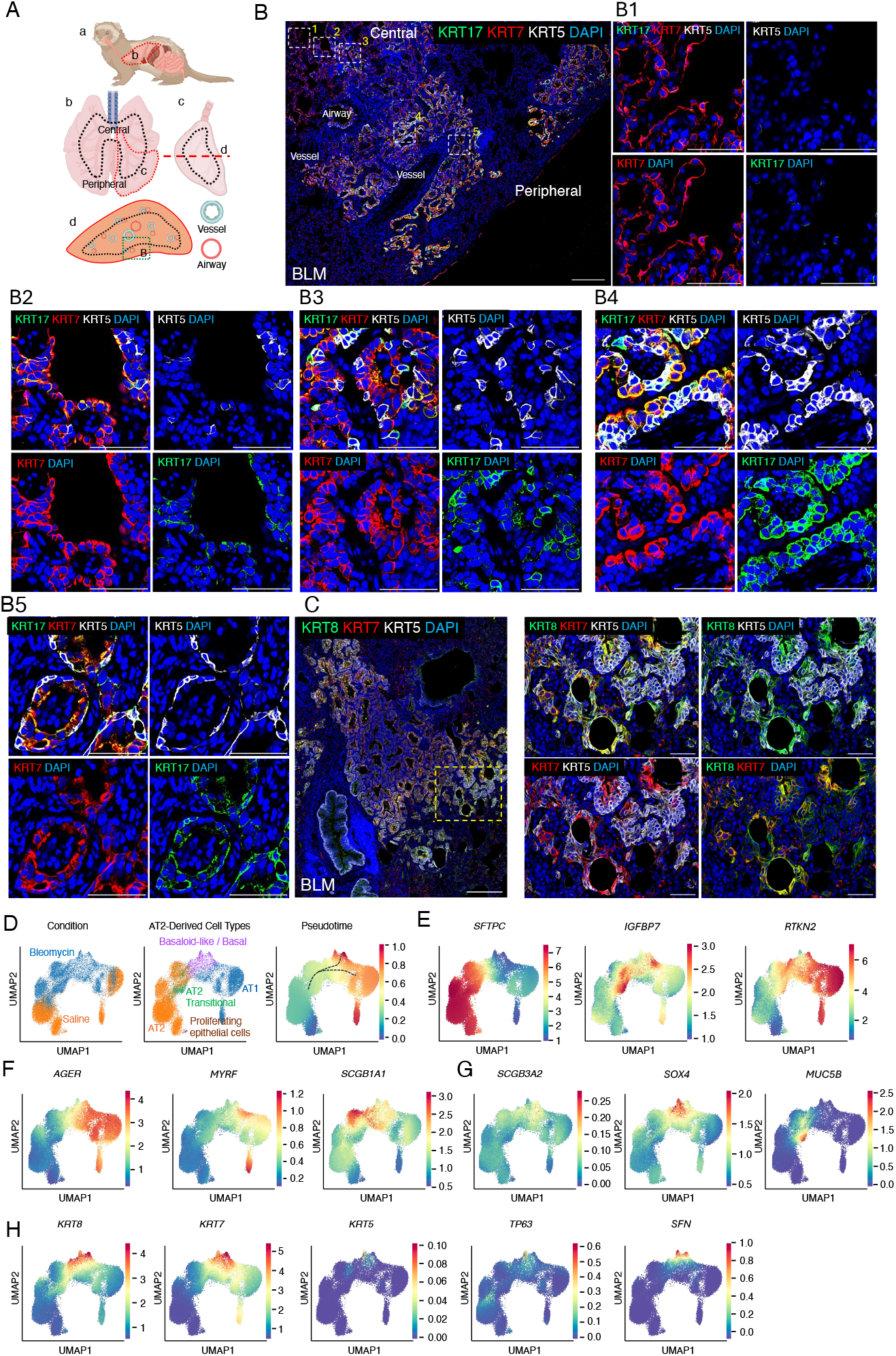
Bleomycin challenge leads to persistence of KRT7^+^ epithelial cells in ferret lung. (A) Schema of ferret internal organs including the lung (a); the central and peripheral locations of all lobes of the lung (b); a single lobe (c) with the level at which cross sections were cut indicated (red dashed line); and a cross section of a lobe (d) used for IF staining and tile imaging from center to the periphery of the lung of in bleomycin-treated ferrets (B). (B) Representative IF images of colocalization of KRT5, KRT7 and KRT17 showing the different states of aberrant epithelial cells from the center to the periphery (1-5) in fibrotic ferret lungs. (B1-B5) Single-channel images of the corresponding boxed areas a-e in B, showing the patterns of expression of epithelial cell markers KRT5, KRT7 and KRT17 from the lobar center to the periphery in fibrotic ferret lungs. (C) Representative IF images of co-staining of KRT8, KRT7 and KRT5. Images in right panels are enlarged double-channel images of the boxed area in the left panel. (D) Pseudotime trajectory analysis of AT2, transitional AT2, Basaloid-like/Basal, and AT1 cells. (E) UMAP plot showing expression of the AT2 marker *SFTPC* along the trajectory. (F) UMAP plot showing expression of the AT1 marker *AGER* along the trajectory. (G) UMAP plots showing expression of the secretory-cell markers *SCGB1A1 and SCGB3A2* along the trajectory. (H) UMAP plots showing expression of the basal-cell markers *KRT7, KRT8, KRT5 and TP63* along the trajectory. Scale bars in B and C equal 200 μm; in B1-B5, and right-hand panels of C they equal

Evidence from human IPF and rodent lung injury models have suggested that AT2 cells progressively transdifferentiates into various aberrant epithelial cell types within fibrotic lesions. Therefore, to track the origin of aberrant epithelial cells in ferret fibrotic lesions we performed pseudotime analysis on AT2 cells. Notably, ferret AT2 cells show multiple distinct trajectories (Fig. 6D, Suppl. Fig. S9), including transitional AT2 cells (defined by SFTPC ^low^, IGFBP7^+^, RTKN^+^, KRT8^low^ expression) to immature AT1 (AGER^+^/MYRF^low^) and then to mature AT1 cells (AGER^+^/MYRF^high^) (Fig. 6E,F and Suppl. Fig. S10A-C). Alternatively, transitional AT2 cells can take a distinct trajectory to KRT8^high^/KRT7^high^/SOX4^+^ and then to KRT8^high^/KRT7^high^/SFN^+^/TP63^+^/KRT5^low^ “basaloid-like” cells (Fig. 6G,H and Suppl. Fig. S10A-C). In addition to these key changes, we also observed changes in additional markers that are described in human IPF and rodent fibrosis scRNA-seq data sets (Suppl. Fig. S10A-C) [33-35, 41, 46, 47]. These observations in conjunction with immunofluorescent localization data suggest that denovo bronchiolization of the distal lung epithelium occurs in bleomycin-induced ferret fibrotic lungs, as postulated in humans [11, 48], and that AT2 cells are a likely progenitor cell intermediate involved in formation of the dysplastic proximalized epithelium in ectopic bronchiolized structures that develop in human IPF lungs.

### Mesenchymal cell dynamics in bleomycin-injured ferret lungs

Focusing on transcriptional alterations in distal lung mesenchymal cells after bleomycin injury, we analyzed 19,628 mesenchymal cells and identified six well segregated mesenchymal cell clusters with unique signature genes (Fig. 7A, B). These included adventitial fibroblasts (*DCN, FBLN1, PI16* and *GLI2*); alveolar fibroblasts (*PDGFRA, TCF21*, and *FN1*); mesothelial cells (*WT1, HAS1* and *SOD3*); myofibroblasts and smooth muscle cells (*ACTA2, CNN1, MYH11, RUNX1, CP* and *TMP1*); pericytes (*PDE5A, PDGFRB* and *POSTN*); and transitional alveolar fibroblasts, which express a combination of fibroblast/pericyte (*PDGFRA, TCF21* and *PDE5A*), myofibroblast (*RUNX1, SFRP1* and *CP*) and collagen (*COL1A1, COL3A1* and *COL5A1*) markers (Fig. 7C and D). These annotations were validated by the clustering and consistent with results from other lung studies [34, 49-52]. Additionally, the analysis of snRNA-seq data revealed a significant increase in the proportion of myofibroblasts and mesothelial cells in response to bleomycin, while there was a decrease in alveolar/lipo fibroblasts (Fig. 7E left panel). This change was accompanied by elevated expression of collagen genes (Fig. 7E right panel). Immunofluorescence staining further showed an increased abundance of CNN1^+^/ACTA2^+^-activated myofibroblasts (Fig. 7F) that increased with higher Ashcroft scores (Suppl. Fig. S11), and accumulation of collagen 1A1 and KRT7^+^ cells surrounding myofibroblast foci (Fig. 7G,I) with interspersed SPC-expressing AT2 cells (Fig. 7H,I) in bleomycin injury compared to control lungs. These observations support the close epithelial-mesenchymal interactions that contribute to the progression of fibrotic lesions and are consistent with findings from the human IPF lung [53]. CellphoneDB analysis of ligand-receptor pairs predicted that KRT7^+^ transitional cells may secrete the ligands TGFB and EREG, which could interact with receptors found in various fibroblast subtypes and other mesenchymal cell types (Fig. 7J). Similarly, IGF1 was expressed by adventitial and transitional fibroblasts, and to a lesser extent by myofibroblasts/smooth muscle cells. The corresponding receptor (IGFR1) was present in both KRT7^+^ transitional cells and AT2 cells, as well as in other mesenchymal cells (Fig. 7J).

**FIGURE 7.**
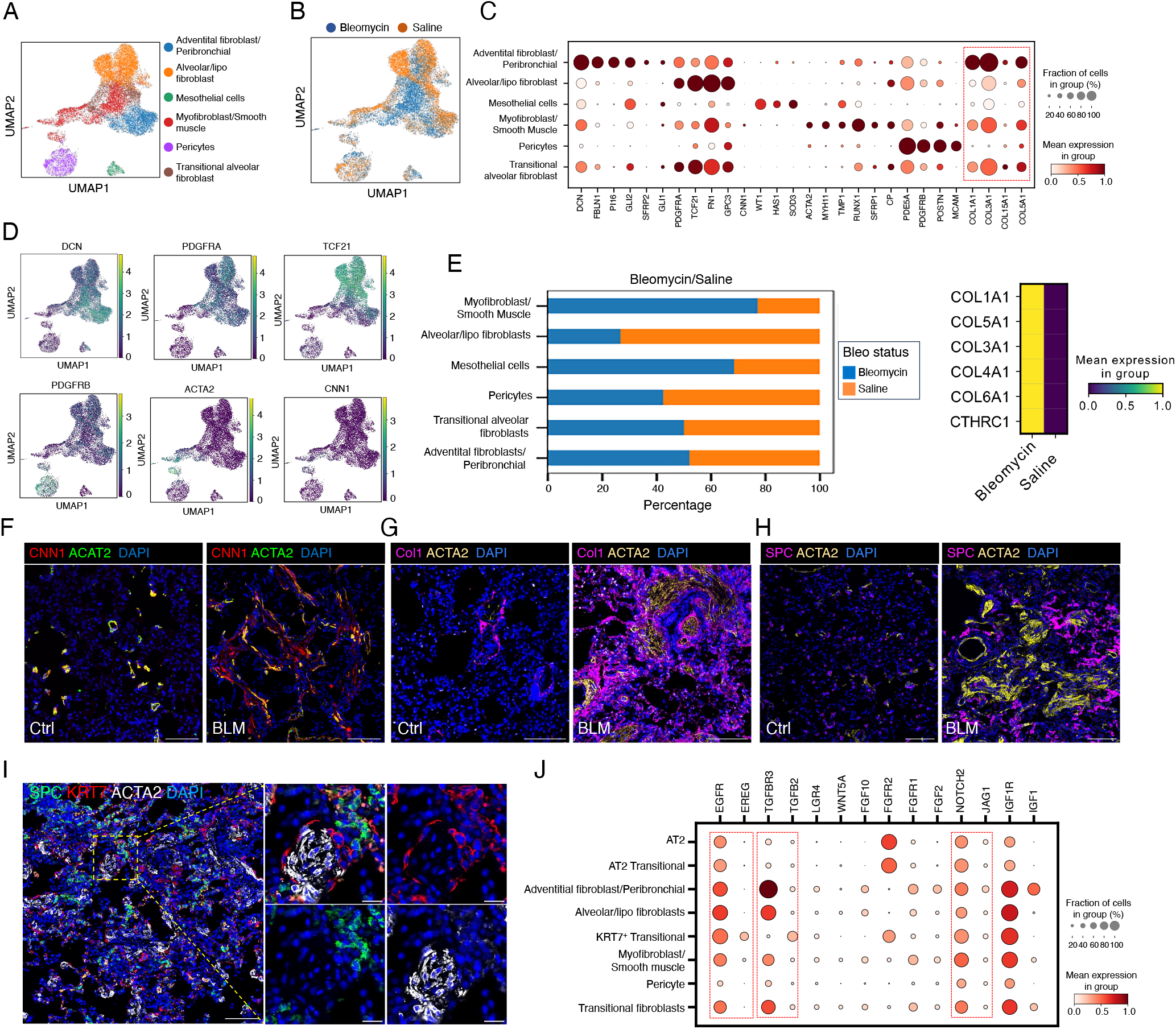
Bleomycin challenge leads to an increase in number of collagen-producing fibroblasts. (A) UMAP showing annotated mesenchymal cell subtypes snRNA-seq data. (B) UMAPs showing differences in types and proportion of mesenchymal cells between ferrets treated with bleomycin and saline. (C) Dot plot showing the frequency of expression of markers of cell types and transitional cell states. (D) UMAPs showing expression of the *DCN, PDGFRA, TCF21, PDGFRB, ACAT2* and *CNN1* genes. (E) (Left panel). Bar plots showing the proportion of each mesenchymal-cell subtype in bleomycin- and saline-treated lungs. (Right panel) Heatmaps of the differential expression of collagen genes in bleomycin- and saline-treated ferret lungs. (F) Representative IF images showing co-staining of ACTA2 (green) and CNN1 (red) in saline control (left panel) and bleomycin-injured (right panel) ferret lungs. (G) Representative IF images of co-staining of Collagen1 and ACTA2 in bleomycin-injured (right panel) and saline control (left panel) lungs. (H) Representative IF image of co-staining of SPC and ACTA2 in saline control (left panel) and bleomycin-treated (right panel) ferret lungs. (J) Representative IF image of co-staining for SPC (green), KRT7 (red), and ACTA2 (white) in bleomycin-treated ferret lung. Inset indicates enlaarged region and individual channels of regions marked by yellow dashed box. DAPI-nuclei (blue). Scale bars in E-I equal 100 µm; in enlarged panels in I, they equal 20 µm. (J) Dot plot showing the ligands and receptor expressed by respective cell types.

## DISCUSSION

Here we present a ferret model of bleomycin-induced acute lung injury, which develops chronic dysplastic regenerative disease resembling human IPF. The key shared histopathological, cellular, and molecular features included fibrotic foci formation, honeycomb cyst-like structures, bronchiolization, and the emergence of aberrant KRT7^+^/KRT17^+^/KRT5**^™^/**TP63^+^ basaloid-like cells [33, 34, 54]. This ferret model also recapitulated the physiologic changes in pulmonary compliance and radiographic imaging observed in individuals with IPF. Due to the anatomical, cellular, and molecular differences between mouse and human lungs, as well as the disease mechanisms in pulmonary fibrosis [55-57], the ferret presents promising opportunities for studying IPF pathophysiology.

A notable difference between humans with IPF and rodent models of lung injury is that the individuals with the condition often develop permanent, chronic, and progressive fibrosis, whereas rodents generally exhibit self-limiting disease progression and reversible fibrosis [4, 5, 58]. However, recent studies using very old mice have shown that more permanent bleomycin-induced fibrosis can also occur in aged rodent models [6]. Our study also showed that younger ferrets had a greater ability to recover pulmonary lung function following bleomycin lung injury. Our chronic state extended to 12 weeks after bleomycin injury; however, another recent ferret study shows permanent fibrotic injury and dysplastic repair to 22 week following bleomycin injury [24]. It remains to be seen if the age dependent irreversible nature of human IPF is different between rodents and ferrets due to differences in lung anatomy and cell biology, such as the presence of RAS/TRB-SC stem cells and respiratory bronchioles, both of which are absent in mice [11, 36, 56].

Our snRNA-seq analysis of saline and bleomycin-treated ferret lungs further identified molecular features reflective of substantial epithelial remodeling following injury and cellular transitional states that mirrored those seen in humans and nonhuman primates [11, 36, 59]. Trajectory analysis of single cell-transcriptomes supports AT2 cells acquire a transitional state that bifurcates to generate both AT1 and basaloid-like cells, a finding similar to human IPF lungs. However, these analyses have not resolved whether RAS/TRB-SC [11, 36] or AT0 cells [11] contribute to dysplastic regenerative states in bleomycin treated lungs.

Recent studies in mice have introduced the concept of injury-induced tissue stem cell niches that form following acute lung injury [60]. This study proposes that injury-induced niches can drive dysplastic nonproductive repair leading to disease pathology and compete with niches that drive euplastic productive repair. Although these proposed mechanisms for IPF in humans are largely based on studies in transgenic mouse models, it remains to be seen if the biology and cell types involved in injury-induced niches and dysplastic repair differ in larger species, like ferrets, and if they bear greater similarity to those seen in humans. Given anatomical and cellular differences in the respiratory zones, precursor populations contributing to proximalization of the airspaces and basaloid cells, which drive structural and functional pathology in pulmonary fibrosis, may vary significantly between rodents and humans [56]. Ferrets may provide an opportunity to bridge this biologic and scientific gap.

Fibrosis-induced proximalization of the airspace typically initiates at the plural surface, or along the interlobular septa, and progresses toward the center of the lobe [42, 43, 61]. Current thinking is that these regions of the lung have unique physical forces and a biochemical makeup rich in collagens that enhances fibrogenic signals in the setting of inflammation and injury [43, 61]. The ferret bleomycin model appears to reproduce this unique aspect of anatomical fibrotic disease progression in IPF.

In summary, we demonstrate that the bleomycin-induced ferret model emulates critical features of IPF, positioning it as a promising alternative to rodent models that fail to replicate certain aspects of its complex pathology. Future studies can leverage this model to explore cellular mechanisms and test IPF therapies, potentially bridging the gap between preclinical findings and human clinical applications.

## MATERIALS AND METHODS

### Creation of an IPF ferret model using bleomycin challenge

All animal experiments were approved by the Institutional Animal Care and Use Committee (IACUC) at the University of Iowa. Male and female domestic ferrets (3–24 months) were obtained from Marshall Farms (North Rose, NY, USA) and housed under controlled temperature (20–22°C) with a light cycle of 16 hours light/8 hours dark and *ad libitum* access to water and diet. Animals were used for experimental purposes after 2 weeks of acclimatization. Ferrets were injured by administering three doses of BLM (2.5 units/ml/kg in saline, Fresenius KABI, Chicago, IL) at 1-week intervals *via* the laryngotracheal route, using a MADgic® Laryngo-Tracheal Mucosal Atomization Device (Teleflex, Morrisville, NC). The first dose of BLM was administered on the first day of Week 0. Throughout the study, all ferrets were monitored daily for activity level, body weight, and respiratory rate.

### Procedure for Anesthesia

Animals were anesthetized by subcutaneous injection of ketamine (5–25 mg/kg) and xylazine (0.5–1.5 mg/kg). Additional isoflurane (0%–5%) supplemented with oxygen flow was provided to sustain anesthesia during pulmonary function testing, CT scanning or intrapulmonary delivery of BLM. Throughout the procedure, temperature, pulse, oxygen saturation, respiratory rate, and general clinical condition were monitored and recorded by trained personnel. Anesthesia was reversed immediately after the procedure was completed by intramuscular injection of atipamezole (0.5 mg/kg). Animals were returned to their cage after they were fully recovered, following a monitoring period.

### Assessment of pulmonary function

In ferrets, PFT was performed using a previously described protocol [19]. In brief, ferrets were anesthetized and intubated with an appropriately sized cuffed endotracheal tube (male I.D.: 3.0 mm, O.D.: 4.2 mm; female I.D.: 2.5 mm, O.D.: 4.0 mm) by a trained operator using a laryngoscope. Intubation was confirmed during a pre-calibration procedure by visualizing the passage of the laryngeal lumen through the cords and examining pressure tracings on a flexiVent ventilator (SCIREQ Inc., Montreal, Quebec, Canada). Routine ventilation parameters were a tidal volume (Vt) of 10 ml/kg, a positive end-expiratory pressure (PEEP) of 3 cmH_2_O, and a respiratory rate of 60 breaths/min (bpm), which resulted in mild hyperventilation. Pulmonary function was assessed using the FX Module 6 of the flexiVent mechanical ventilator system, a computer-controlled piston ventilator that allows control of ventilation parameters [15]. Anesthetized ferrets were placed on the heated plate, and the endotracheal tube was connected to the in-line ventilator. Baseline parameters were calibrated and acquired, and then multiple forced maneuvers were performed to assess parameters of pulmonary functions, including inspiratory capacity (IC), forced expiratory volume (FEVx) at defined times (defined as “x” seconds), forced vital capacity (FVC), resistance of the respiratory system (Rrs), elastance of the respiratory capacity (Ers), compliance of the respiratory system (Crs), Newtonian resistance (Rn), tissue damping (G), tissue elastance (H), pressure volume (PV) loop, and quasi-static compliance (Cst). For flexiVent perturbations, a coefficient of determination of 0.95 or above was considered acceptable. All test maneuvers and perturbations were conducted until three acceptable measurements were collected. The collected data were analyzed using the pre-installed Scireq flexiWare Version 8.0, Service Pack 4 (Montreal, QC Canada). For each mechanical parameter, a total of six technical replicate measurements were performed and the average value was used for data representation.

### CT imaging and quantification

Ferrets were anesthetized and imaged in the prone position using a SOMATOM Force dual-source CT scanner (Siemens Healthineers, Germany) in the Flash (dual-source) scanning mode. The ferrets breathed spontaneously during scanning. Full inspiration (end-tidal: ET) scans were obtained by scanning each ferret continuously 5 to 10 times and selecting the image with the largest lung volume as the ET scan. All images were acquired with the following parameters: detector configuration, 192 × 0.6 mm; kVp, 100; mAs: 100; slice thickness, 0.6 mm; slice interval, 0.3 mm; reconstruction kernel, Qr49 ADMIRE 3; scan pitch, 2.8; rotation time, 0.25 seconds. Quantification was performed using the Pulmonary Analysis Software Suite (PASS) (University of Iowa, Iowa City, Iowa), a software package that was developed in-house and customized for volumetric analysis of lung opacification by Hounsfield units (HU) in ferret lung disease models [62]. The left and right lungs were each segmented first, and then histogram analysis was applied to each segment to identify the high-density (fibrotic-like) lung regions based on the percentage of lung voxels with intensity values falling between -500 HU and 0 HU [62, 63].

### Analysis of histopathology

Ferrets were euthanized when in clinical distress or at the end of the study. All five main lobes (except the accessory cordate lobe) were separately isolated. The middle portion of each lobe was fixed with 10% neutral-buffered formalin (10% NBF) and embedded for histopathological and immunohistochemical (IHC) analyses. The remaining lung tissue was snap frozen in liquid nitrogen (LN_2_). Collagen deposition in lung was semi-quantified based on Masson’s trichrome staining of sections [64]. Morphological changes of fibrosis in hematoxylin and eosin (H&E)-stained lung sections were graded using the Ashcroft scale, by three independent researchers blinded to experimental conditions [65, 66]. For this analysis, all parenchyma in the lung sections were assessed at 5x magnification after slide digitalization, using the 0–8 point Ashcroft scale. The mean of three individual scores of all five lobes served as the final Ashcroft score for analysis of each animal [65, 66].

### Measurement of hydroxyproline content

Plasma and tissue from the combined lung lobes were homogenized for isolation of total protein. The HYP content of each sample was measured using a hydroxyproline assay kit (Ab222941, Abcam, Waltham, MA) according to the manufacturer’s instructions. The HYP content of plasma is presented as µg/mL of volume, and that of wet lung tissue as µg/mg of protein.

### Immunofluorescence (IF) staining

Frozen sections (8-µm) of lung tissues embedded in optimal cutting temperature (OCT) compound were air dried for 30 minutes at room temperature before being post-fixed in 4% paraformaldehyde (PFA) for 10 minutes. Tissue sections were then permeabilized in 0.2% Triton X-100 in PBS for 30 minutes, blocked with 5% donkey serum for 2 hours, incubated with primary antibody in diluent buffer (1% donkey serum, 0.03%Triton X-100, and 1 mM CaCl2 in PBS) at 4°C overnight, and incubated with Alexa Fluor-labeled secondary antibody at room temperature for 2 hours. Paraffin sections were prepared similarly, but were subjected to deparaffinization, rehydration, and antigen retrieved, before being blocked in blocking buffer. Slides containing nuclei that were stained using Hoechst 333427 (Invitrogen, Carlsbad, CA, USA) were then mounted with VECTASHIELD Antifade Mounting Medium (Vector Laboratories, Burlingame, CA, USA) and imaged on a Zeiss LSM880 confocal microscope (Carl Zeiss Meditec AG, Jena, Germany). All images were processed using FIJI-ImageJ software. The primary and secondary antibodies used for IHC staining are listed in Suppl. Tables S1 and S2.

### Measurement of proteins in plasma using in-house enzyme-linked immunosorbent assay (ELISA)

Ferret plasma was diluted (1:10) in carbonate-bicarbonate buffer. Diluted plasma (200 µl) was used to coat each well of a polyvinyl chloride (PVC) microtiter plate at 4°C overnight. The coated wells were rinsed three times with washing buffer (PBS and 0.05% Tween 20 (pH 7.4)) and blocked with 5% non-fat milk in washing buffer for 2 hours at 37 °C. After washing three times for 5 minutes each, 100 µl of diluted primary antibody (1:1000 in washing buffer) (Suppl. Tables S1 and S2) for the protein of interest was added to the wells and incubated at 37°C for 2 hours. The wells were washed five times for 5 minutes each and then incubated with HRP-conjugated secondary antibody (1:2000 in washing buffer). After the wells were again washed five times for 5 minutes each, 100 µl of TMB substrate solution (Sigma, St. Louis, MO) was added to each and the samples were incubated in the dark for 15–30 at room temperature, at which point 100 µl of stopping solution (2M H_2_SO_4_) was added to the wells. Optical density (OD) of the xx plate wells was measured at 450 nm (OD_450nm_) using a SpectraMax i3x Microplate Reader (Molecular Device, San Jose, CA). A no-plasma (PBS alone) well served as a blank control. The results are presented as ΔOD_450nm_, defined as the difference between the value of the OD_450nm_ readout of the experimental sample substrate and the readout of the blank control.

### Isolation and 10x sequencing of single nuclei RNA from ferret lung

Lung nuclei were isolated using a CST buffer (20 mM Tris-HCl pH 7.5, 292 mM NaCl, 2 mM CaCl2, 42 mM MgCl2, 1% CHAPS, and 2% BSA) as described previously [67]. Briefly, snap frozen lung tissues (50 mg) were immediately placed into a nuclei isolation using CST buffer, and a dounce homogenizer was used to mechanically dissociate nuclei from cells. The resulting homogenate was filtered through a 40 µm strainer, and nuclei were pelleted by centrifugation at 300 × g for 2 minutes at 4°C. Nuclei were resuspended in 0.04% BSA buffer, to a final concentration of 1,000 nuclei/µl. Single-nuclei RNA sequencing (snRNA-seq) libraries were prepared using the 10x Genomics 5’ Kit v2, following the manufacturer’s protocol (10X Genomics). Libraries were sequenced on an Illumina Nextseq 550 platform using 75-cycle paired-end sequencing, and 8bp for the index read, 28bp for the R1 read, and 56bp for the R2 read.

### Processing and analysis of snRNA-seq data

Illumina BCL files were processed to fastq format using the bcl2fastq v2.2.0 conversion software. To generate a count matrix, the sequenced files from each independent sample were processed through 10X the Genomics Cell Ranger software v7.1.0, using the default “include intron” mode and a reference build from Mustela putorius furo (domestic ferret) reference genome ASM1176430v1.1.

Samples were annotated for ambient content using Dropkick (https://github.com/KenLauLab/dropkick) and annotated with doublet scores using Scrublet (https://github.com/swolock/scrublet), both of which are built into the Scanpy toolkit (https://github.com/scverse/scanpy). Filtered feature-barcode matrices from Cell Ranger were initially filtered using Cell Ranger knee cutoffs to remove cells with unique molecular identifier (UMI) counts above the 99 and below the 4 percentile, unless the 4th percentile UMI count was less than 800, in which case a floor of 800 UMI was used. Cells annotated as doublets, cells with greater than 4% mitochondrial UMIs, and cells with a Dropkick score below 0.25 were then removed.

### Visualization, dimensionality reduction, and clustering of raw data

The filtered raw count data were then annotated using a reference-based annotation tool (https://github.com/gordian-biotechnology/celltype_predict_ACTINN) using multiple published lung datasets (GSE135893,GSE136831, and the human cell lung atlas). snRNA-seq datasets were normalized and transformed the filtered count matrix as per standard Pearson residuals Scanpy pipeline. Dimension reduction was performed using the top 3,500 highly variable genes, and the top 50 principal components were calculated and used for nearest neighborhood graph Leiden clustering and UMAP visualization. Cell type labels were assigned to Leiden clusters based on consensus cell-type labels from references and specific cell-type markers such as KRT7 to annotate specific subpopulations. All details of data processing, filtering, and annotation are shown in the accompanying Jupyter notebooks.

### Pseudotime analysis

Psuedotime analysis was computed using the Palantir (https://github.com/dpeerlab/Palantir) v1.3.3 software [68, 69].

### Prediction and visualization of cell-cell interactions

Cell-cell interaction scores were computed using the Cellphone database (https://github.com/ventolab/CellphoneDB) v5.0.1 [68, 69].

### Statistical analysis and reproducibility

Data analysis was performed using the GraphPad Prism 9 software (GraphPad Software, San Diego, CA, USA). For all experiments, data are shown as mean ± SEM, unless indicated otherwise. Comparisons between two groups were done using an unpaired two-tailed t-test, and comparisons of more than two groups were analyzed using one-way ANOVA. Figures and legends provide the number of independent experiments and the results of representative experiments.

### Reporting summary

Further information on research design is available in the Nature Portfolio Reporting Summary linked to this article.

## Data availability

snRNA-Seq data were deposited in the NCBI’s Gene Expression Omnibus (GEO) database (GSE279470). Reviewers can access the deposited sequence data at https://www.ncbi.nlm.nih.gov/geo/query/acc.cgi?acc=GSE279470) using the following token (snizcegudxwzbmt). Values for all data points in graphs can be found in the Supplemental Data file.

## Code availability

Themajority of the analysis was carried out using published and freely available software and pre-existing packages mentioned in the methods. No customcodewas generated. R scripts used to analyze data and generate figures are available upon request to VS and XL.

## Author contributions

XL, VS and JFE designed research studies. SW, ML, AT, JA, LY, JW and JM conducted experiments. ID, DP and VS performed snRNA sequencing and analysis. SW, JG, VS and XL analyzed data. SW, VS and XL wrote the manuscript. EAH, YW, PRT and JFE revised the manuscript.

## Acknowledgments

This work was funded by the following grants from the National Institutes of Health NHLBI (Federal Contract 75N92024R00008 and R01 HL165404) to JFE and from NHLBI/NIH (R01HL160939 and R01HL153375) to PRT.

## Ethics approval statement

This study was performed according to protocols approved by the Institutional Animal Care and Use Committee (IACUC) of the University of Iowa and conforms to National Institutes of Health (NIH) standards (Protocol ID1071945). Humane/euthanasia endpoints: Labored breathing with respiratory rate greater than 90 bpm or lack of movement or response upon stimulation.

## SUPPLEMENTARY FILES

### Supplementary Materials

1. Supplemental figures

2. Supplemental tables

