## Supplemental figures and tables for "Ferret model of bleomycin-induced lung injury shares features of human idiopathic pulmonary fibrosis"

**SUPPLEMENTARY FILES**

1. Supplemental figures
2. Supplemental tables

### Supplementary Figures

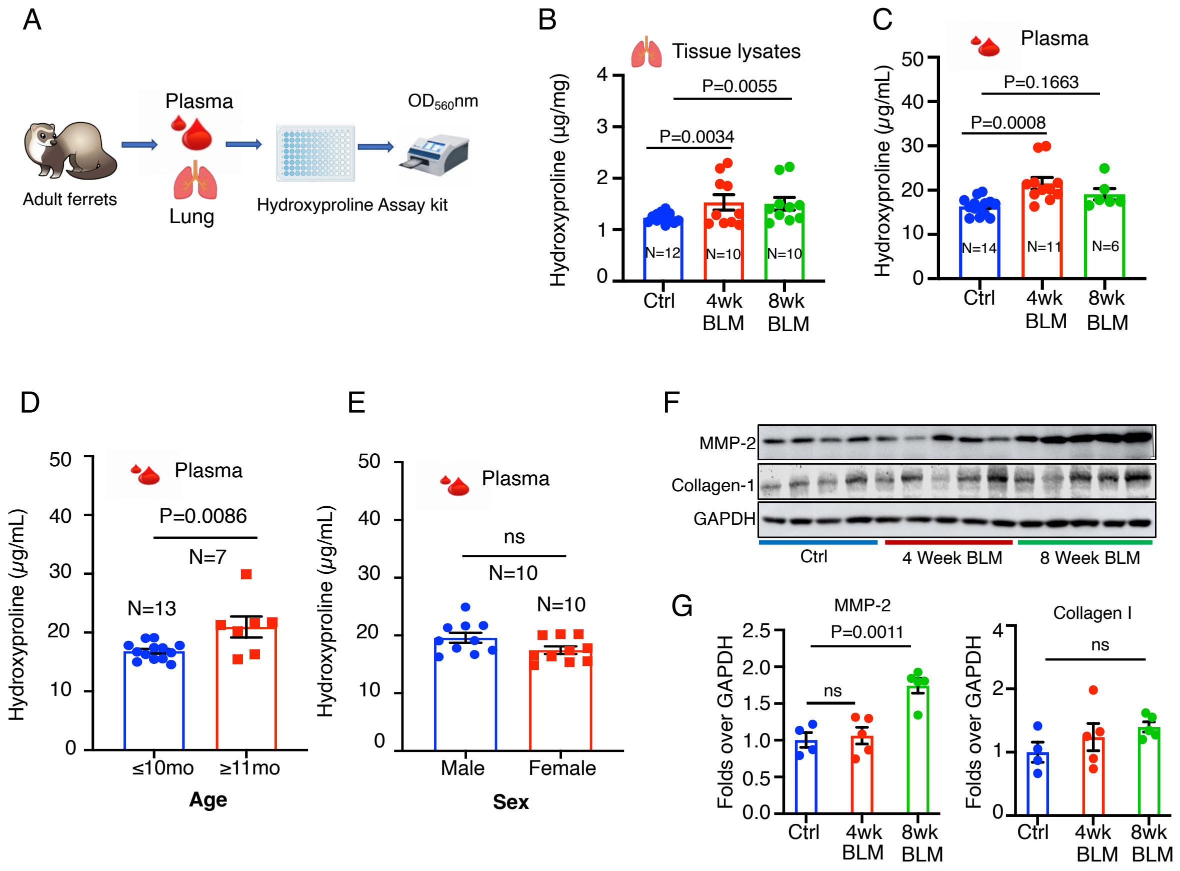

**Figure S1. Hydroxyproline content and extracellular matrix (ECM) deposition are elevated in bleomycin-treated ferrets.** (A) Schema showing the workflow to test circulatory hydroxyproline (HYP) content in the plasma and lung. (B, C) The HYP content in ferret lung tissue (B) and plasma (C) at 4 and 8 weeks after the first dose of bleomycin, compared to those in age- and sex-matched saline controls. (D,E) The HYP content in older (≥11 months) vs. younger (≤10 months) (D) male and female ferrets (E) challenged with bleomycin (P=0.0086, N=7). (F) Representatives immunoblots of matrix metalloproteinase-2 (MMP-2) and collagen-I in the lungs of ferrets at 4- and 8-weeks post bleomycin challenge. (G) Densitometry-based semi-quantification of relative abundance of MMP-2 and collagen-I in bleomycin-injured ferrets over time (P=0.0011, N=5). Data in B, C, and G represent the mean ± SEM and were analyzed by One-way ANOVA, followed by Dunnett's comparison test. Data in D, E represent the mean ± SEM and were analyzed by unpaired *t*-test. Data from individual ferrets are represented as distinct data points in graphs.

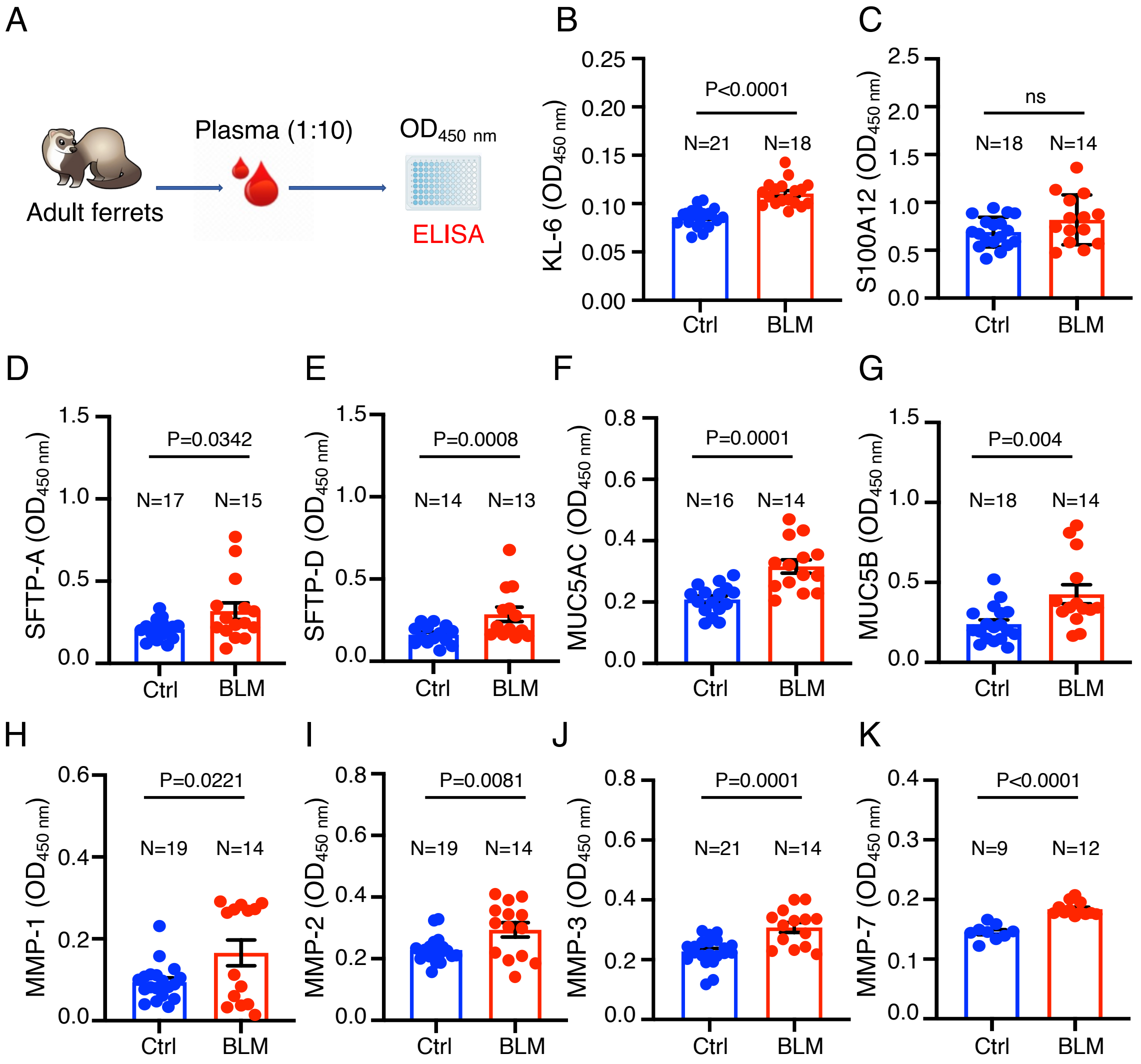

**Figure S2. Circulatory biomarkers of pulmonary fibrosis are elevated in the plasma of bleomycin ferrets.** (A) Schema showing the workflow of in-house developed enzyme-linked immunoassay (ELISA) for assessing expression of biomarkers of PF in ferret plasma samples. Plasma from age- and sex-matched saline-treated ferrets served as controls (Ctrl). (B-K) Comparisons of the abundance of proteins in the plasma of saline- and bleomycin-treated controls included Krebs von den Lungen 6 (KL-6) ($p<0.0001$) (B), S100 calcium-binding protein A12 (S100A12) ($p=0.0910$) (C), surfactant protein A (SFTPA) ($p=0.0342$) (D), SFTPD ($p=0.0008$) (E), MUC5AC ($p=0.0001$) (F), MUC5B ($p=0.004$) (G), matrix metalloproteinase 1 (MMP-1) ($p=0.0221$) (H), MMP-2 ($p=0.0081$) (I), MMP-3 ($p=0.0001$) (J) and MMP-7 ($p<0.0001$) (K) (N, number of ferrets showed in graph). Data represent the mean ± SEM, and GraphPad prism was used to run the unpaired *t*-test. Individual ferret data are represented by distinct data points in the graphs.

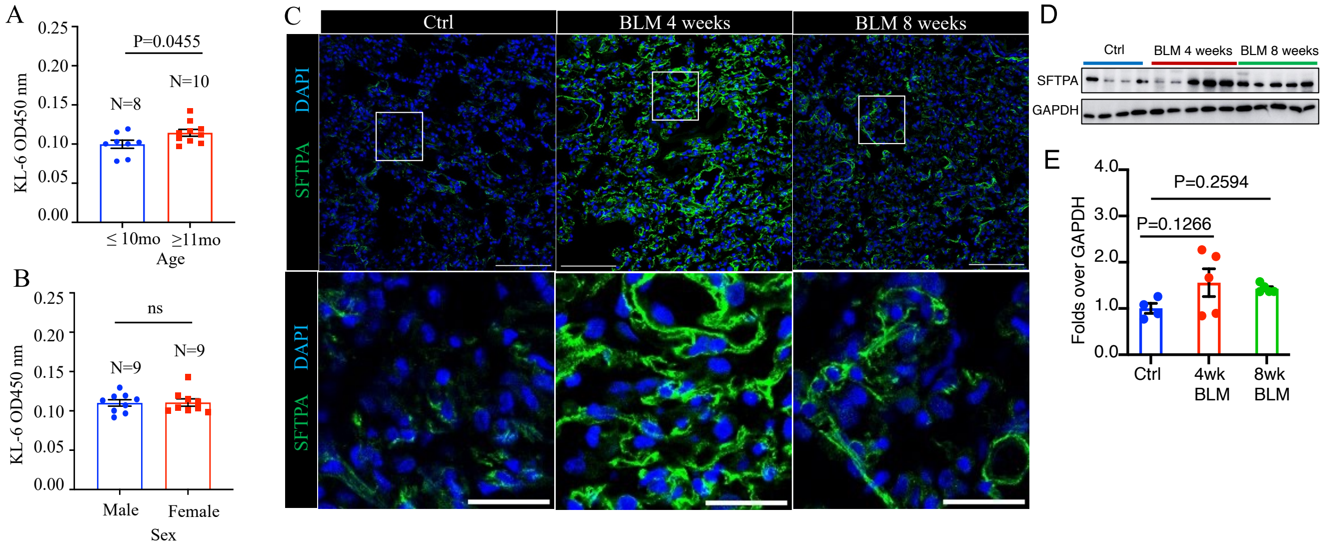

**Figure S3. Bleomycin challenge in ferrets elevates plasma KL-6 and lung SFTPA levels.** (A, B). Comparison of KL-6 levels in the plasma of bleomycin-treated ferrets by age (≤10 months vs. ≥11 months old) (A) and sex (B). (C) Representative IF images of SFTPA in lungs of saline vs. bleomycin-treated ferrets at 4 and 8 weeks post treatment. Bottom panels are enlargements of boxed area of corresponding images in the top panels. (D) Representative Western blots of SFTPA expression in lungs of ferrets at 4- and 8-week post bleomycin challenge. (E) Semi-quantitation of relative abundance of SFTPA protein, assessed by densitometry analysis of Western blots shown in D. Data in A and B are shown as mean ± SEM and were statistically analyzed using the unpaired *t*-test (N=8 for control and 10 for bleomycin in A; N=9 in B); data in E represent the mean ± SEM and were analyzed by One-way ANOVA, followed by Dunnett's comparison test (N=5). Scale bars in top panels represent100 µm; those in bottom panels represent 20 µm. Data from individual ferrets are represented as distinct data points in graphs.

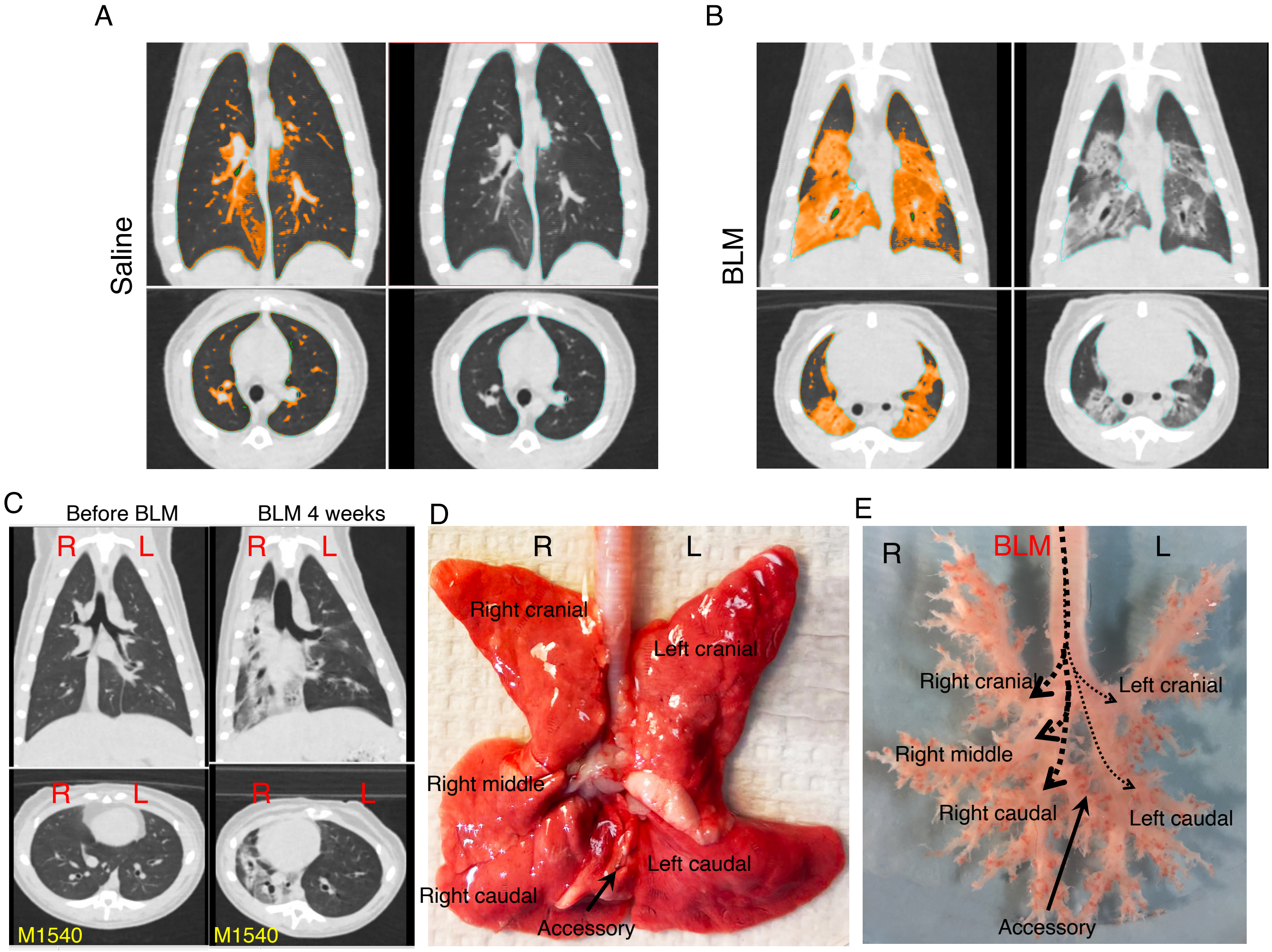

**Figure S4. Radiographic quantification of bleomycin-induced fibrosis in ferret lungs.** Volumetric analysis of CT scans. (A,B) Representative 2D scanning images of ferret lungs treated with saline (A) and bleomycin (B) for HAA quantification. (C) Representative 2D scanning images of lungs of a ferret before (left panel) and 4 weeks after bleomycin exposure, showing that more HAA is present in right vs. left lung (animal ID# M1540). (D) Image of ferret lung with the 5 main lobes (right 3 and left 2), and an accessory lobe. (E) Image of ferret cartilaginous airway tree, showing the potential efficient delivery of agents into different lobes of ferret lung using the intratracheal approach. R, right, L, left.

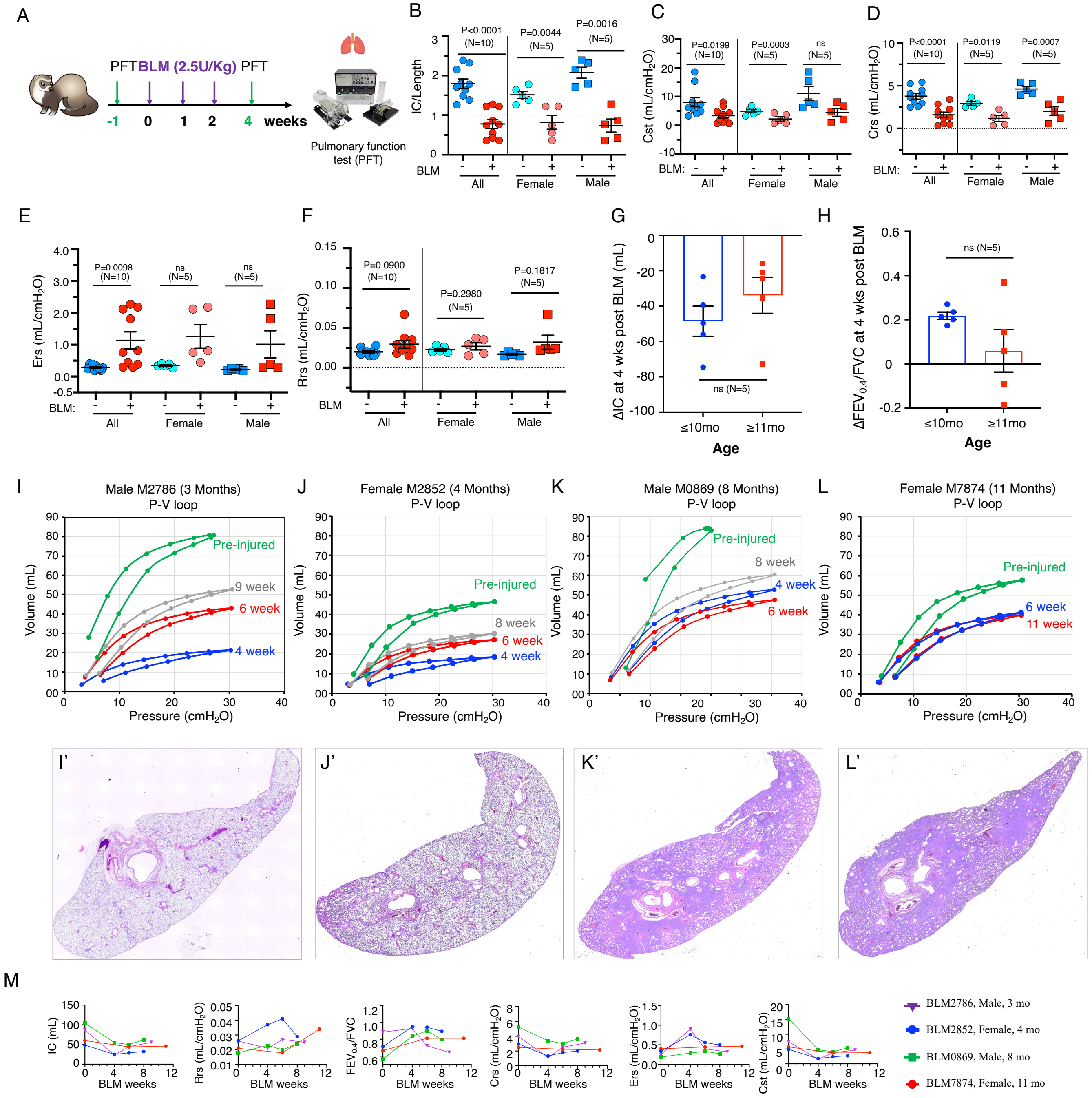

**Figure S5. Bleomycin challenge in ferrets decreases pulmonary compliance.** (A) Schema showing the workflow of the pulmonary function test (PFT) in bleomycin-injured ferrets. The PFT was used at one week before (-1 week) and 4, and/or 6 weeks after the first dose of bleomycin was delivered, for longitudinal monitoring of the PF phenotype in surviving ferrets. The green arrows denote the time that measurements were made. (B-F) PFT metrics include the ratio of IC/length (B), quasi-static compliance (ml/cmH$_2$O) (Cst) (C), dynamic compliance of the respiratory system (Crs, ml/cm$H_2O$) (D), elastance of the respiratory system (Ers, cm$H_2O$/ml) (E), and respiratory system resistance (Rrs) (F), and these were assessed in 10 animals (5 males and 5 females) at one week before and 4 weeks after the first dose of bleomycin was given. (G, H) Changes in IC (G) and $FEV_{0.4}$/FVC ratio (H) at 4 weeks post bleomycin challenge between ages (≤10 months vs. >10 months) (G). (I-L) Changes of pressure-volume loops (PV-loops) as PF developed in 4 ferrets at the indicated ages: 3 months (I), 4 months (J), 8 months (K) and 11 (L) months after the first dose of bleomycin. (I'-L') Representative images of H&E staining in sections of whole lobe from the corresponding ferrets in I-L. (M) Other PFT metrics assessed during the course of development of PF in the above 4 ferrets, including IC, Rrs, $FEV_{0.4}$/FVC ratio, Crs, and Cst. Data in B-F were analyzed for the mixed-sex group ("All" at the left part of each graph, N=10), "Female" (middle part of each graph, N=5), and "Male" (right part of each graph, N=5). All graphs show the mean ± SEM and were analyzed using the unpaired *t*-test. Data from individual ferrets are presented as distinct data points in graphs in B-H.

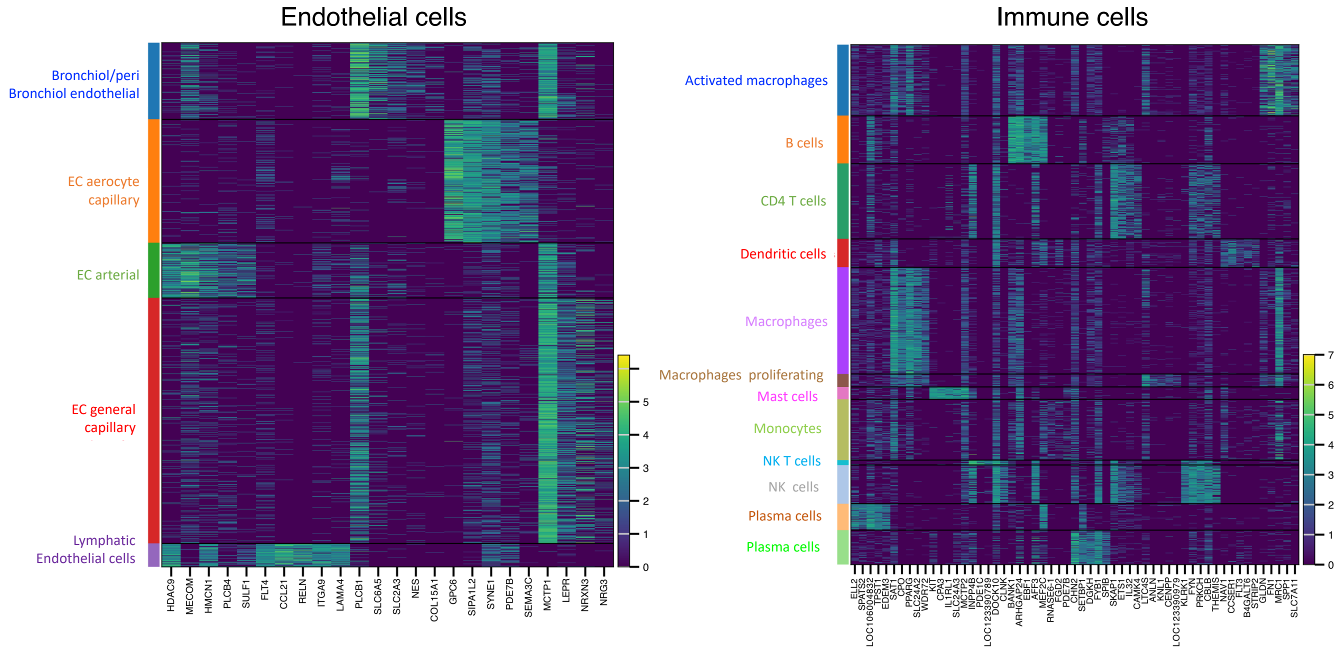

**Figure S6. Single nucleus RNA sequencing (snRNA-Seq) data of ferret lungs.**

(A,B) Heatmaps of the top marker genes in endothelial cell clusters (A) and immune cell clusters (B).

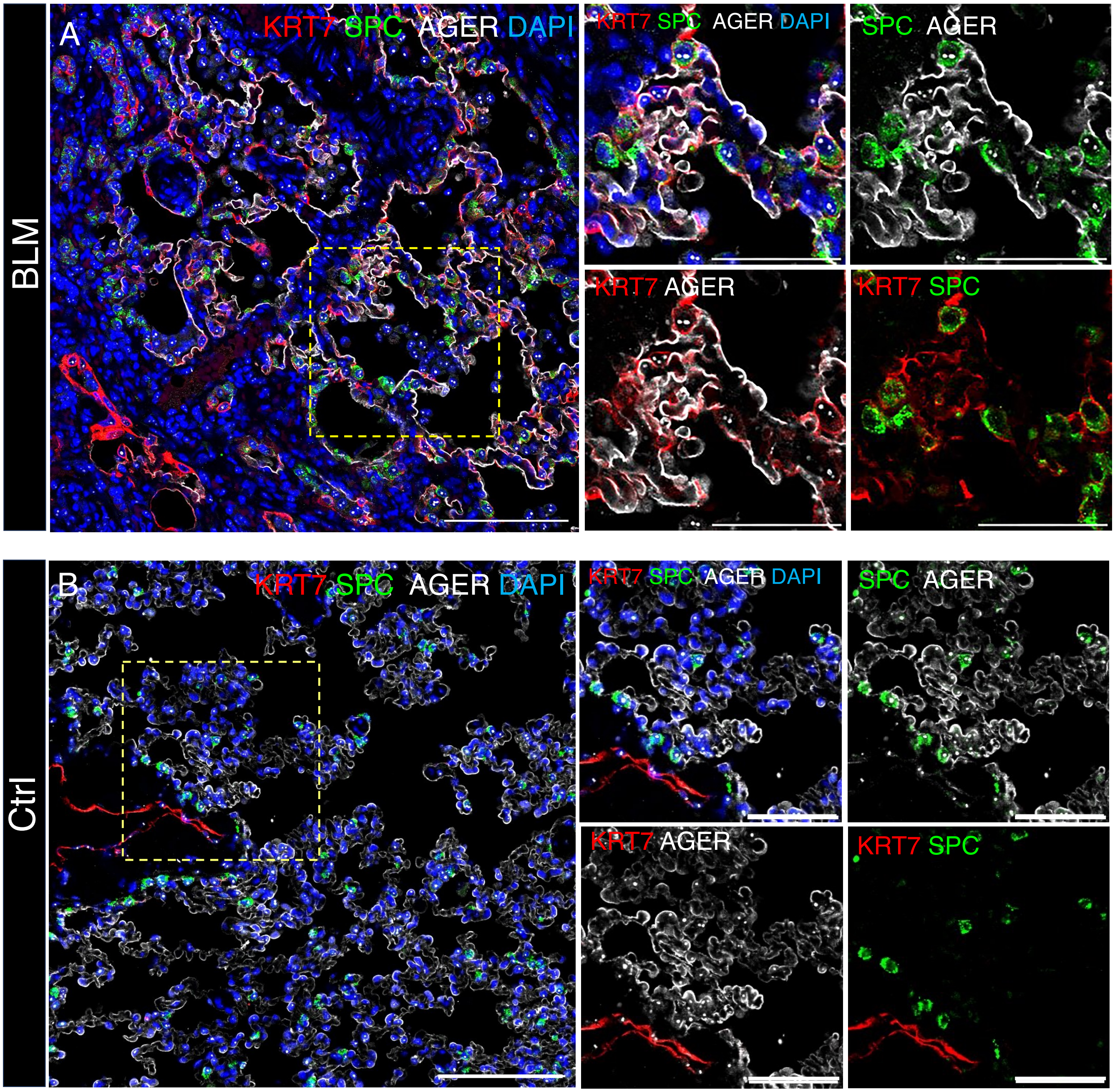

**Figure S7. Bleomycin challenge of ferret results in aberrant KRT7 epithelial cells in the distal lung.**

(A) Representative IF images of co-staining for SPC, KRT7, and AT1 cell marker AGER in bleomycin-induced PF ferret lung. (B) Representative IF images of co-staining for SPC, KRT7 and the AT1 cell marker AGER, in saline control ferret lung. Images in the panels to the right of A-B are enlargements of single- (A) and double-channel images of the boxed areas in the panels at the left. In all images, scale bars equal 50 µm.

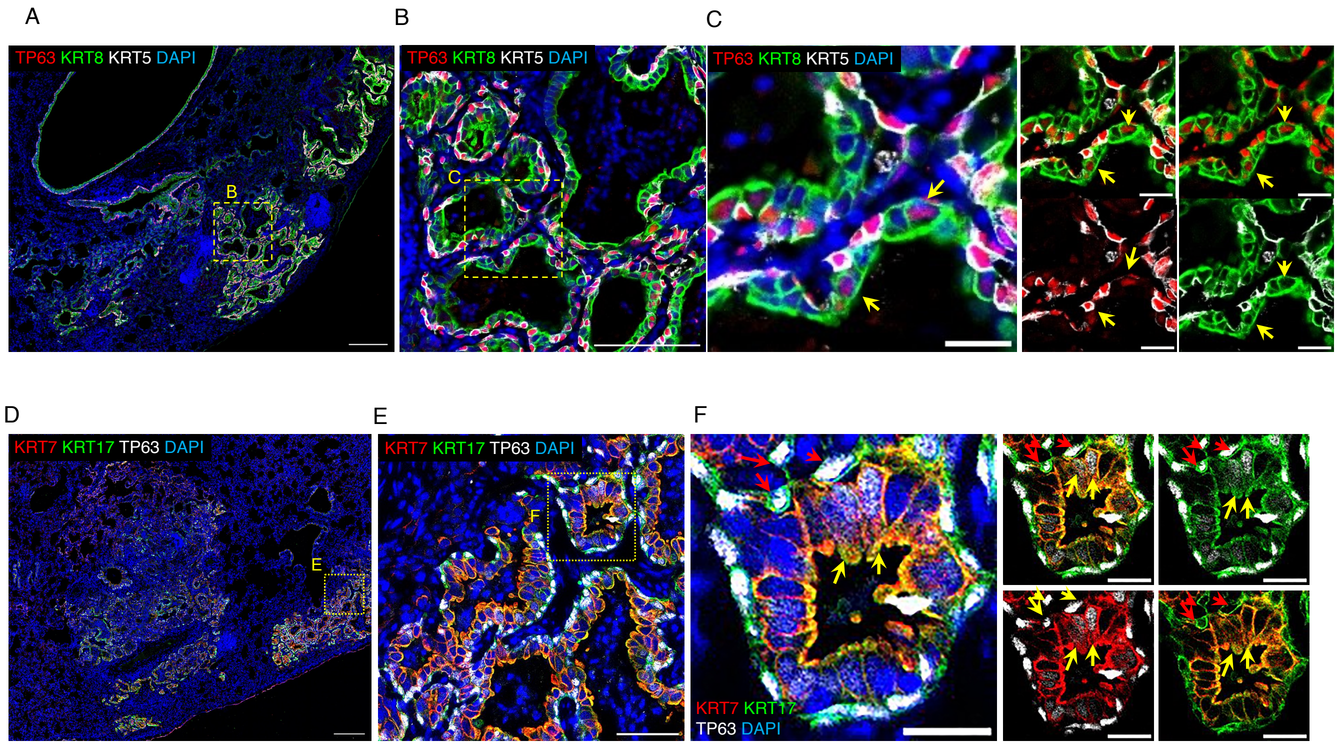

**Figure S8. Implication of KRT7$^+$ epithelial cells in the bronchiolization of bleomycin ferret lungs.** (A) Representative IF image of colocalization of KRT8, KRT5 and TP63. (B) Enlarged image of the boxed area in (A). (C) Enlarged image of the boxed area in (B), showing that most TP63$^+$ cells were colocalized with KRT8$^+$/KRT5$^-$ cells (arrows). Right panels show double-channel images for C. (D) Representative IF image of colocalization of KRT7, KRT17 and TP63. (E) Enlargement of the image of the boxed area in (D). (F) Enlargement of the image of the boxed area in (E), showing that most of TP63$^+$ cells colocalized with KRT8$^+$ cells (red arrows in F), subset of KRT7/KRT17 double-positive cells were TP63$^+$ cells (yellow arrow in F). Right panels, single- and double-channel images of F. Scale bars in A and C equal 100 µm; B and E equal 50 µm; C and F equal 20 µm.

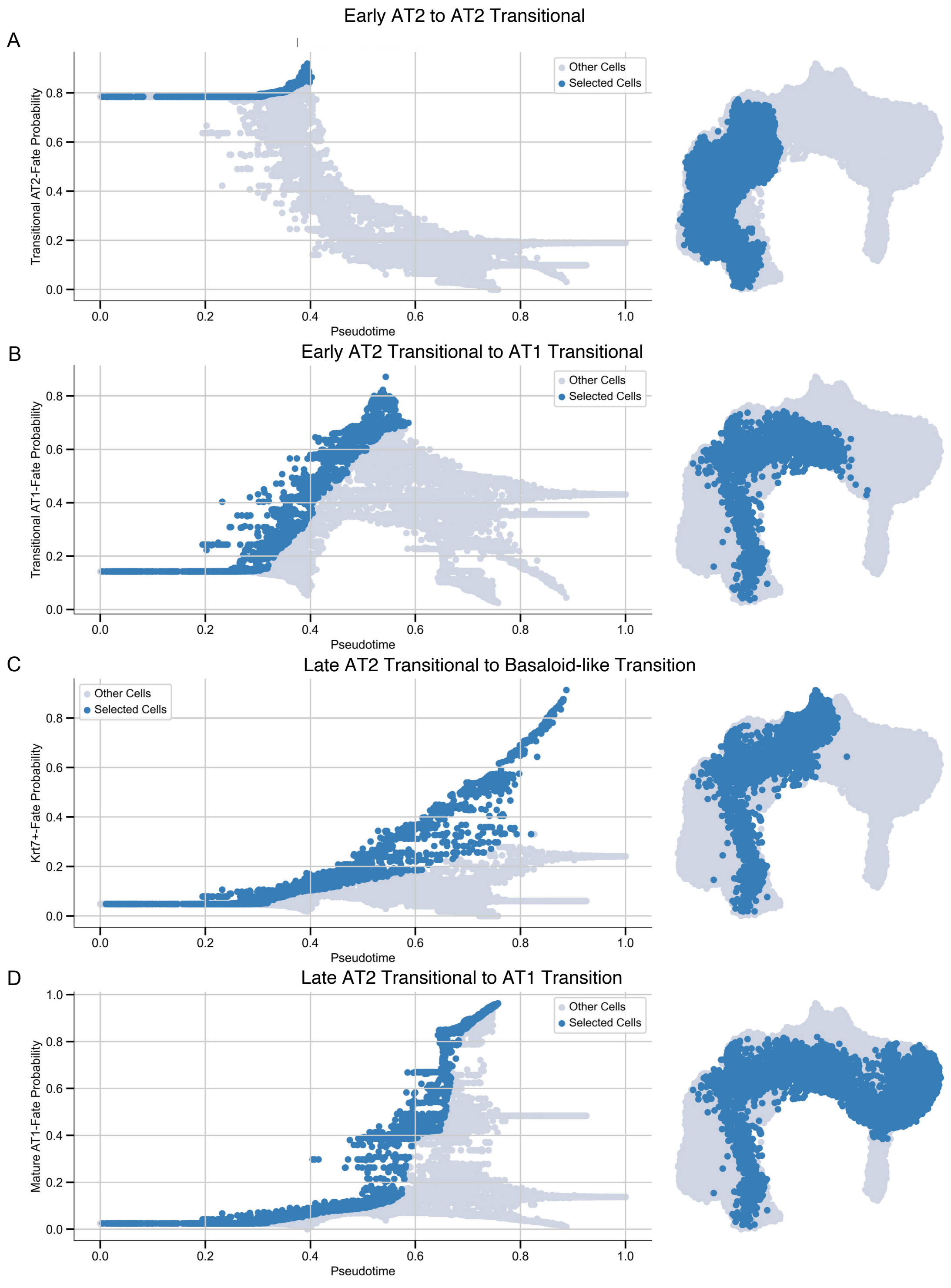

**Figure S9. Trajectory of AT2 cells in bleomycin induced lung fibrosis in ferret.** Using palantir plot_branch_selection function we calculated the fate probability for each cell type versus Pseudotime, Early AT2 to AT2 transitional (A), Early AT2 transitional to AT1 transitional (B), Late AT2 transitional to "basaloid-like" cells transition (C), Late AT2 transitional to AT1 transition (D).

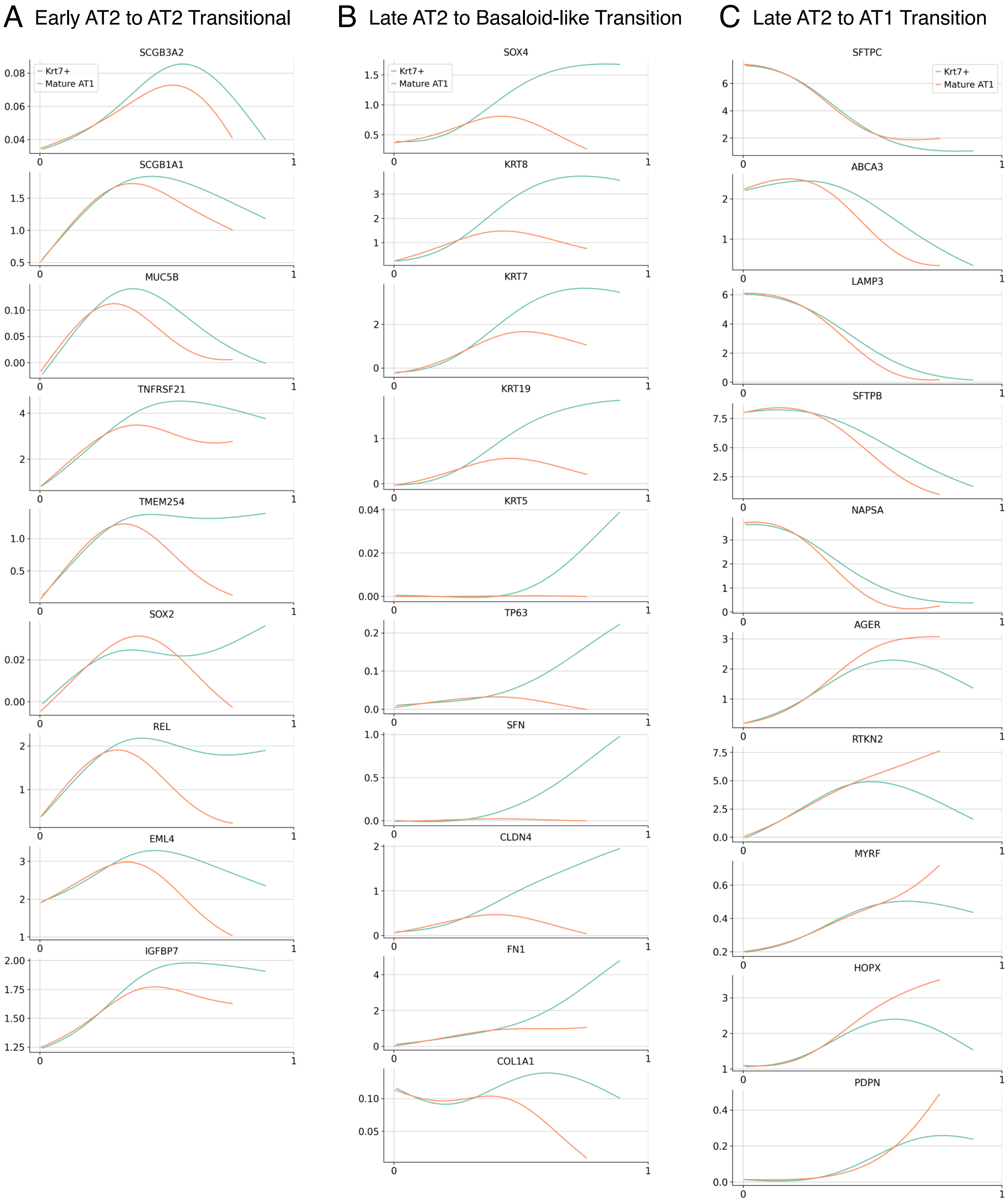

**Figure S10. Changes in gene expression along the trajectory line graph.** (A-C) Line plots representing Early AT2 cell to AT2 transitional cell markers (A), Late AT2 to “Basaloid-like” Transition cells (B), Late AT2 to AT1 cell transitioning marker (C). Y-axis indicates MAGIC imputed gene expression and X-axis indicates pseudotime.

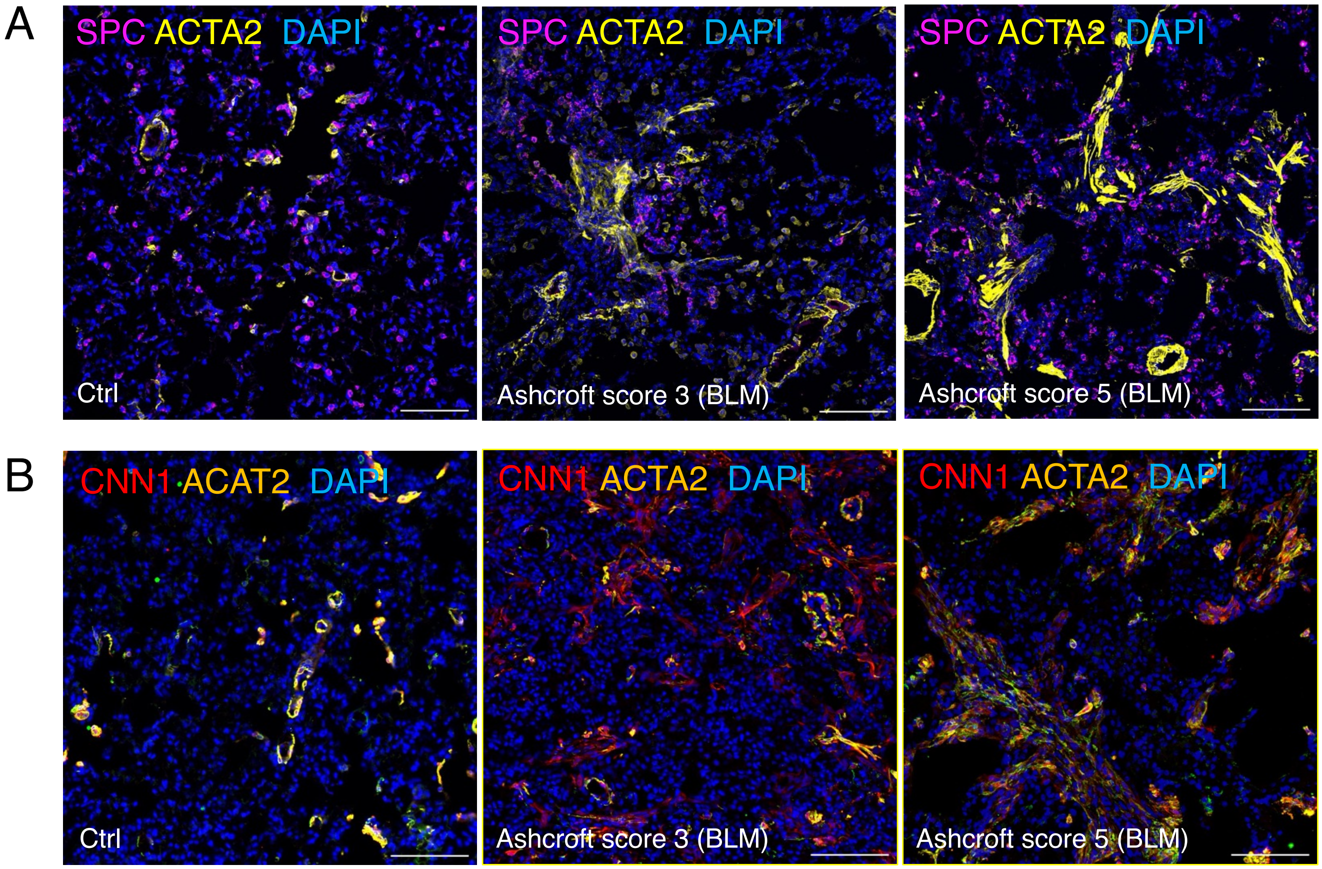

**Figure S11. Bleomycin challenge in ferrets promotes differentiation of myofibroblasts and disease progression in the distal lung.** (A) Representative IF images of colocalization of SPC and ACTA2 in saline control (left panel) and bleomycin-induced PF ferret lung with an Ashcroft score 3 (middle panel) and an Ashcroft score of 5 (right panel). (B) Representative IF images of colocalization of mesenchymal cell marker CNN1 and ACTA2 in saline control (left panel) and bleomycin-induced IPF ferret lungs with an Ashcroft score of 3 (middle panel) and an Ashcroft score 5 (right panel). In all images, scale bars equal 100 µm.

### Supplementary Tables

**Suppl. Table S1. List of primary antibodies**

| Antigen | Vendor | Cat # | Host | Application(s) |
|---|---|---|---|---|
| AGER | R&D systems a biotechne brand | AF1145 | Goat | WB, IHC(P), ELISA |
| α-SMA | proteintech | 14395-1-AP | Rabbit | IF, IHC, WB |
| Collagen III | Aviva | OASB02742 | Goat | FC, IF, IHC, IP, WB, ELISA |
| Collagen 1A1 | Novusbio | NB600-408 | Rabbit | IF, WB, IHC |
| Cytokeratin 5 | Biolegend | 905901 | Chicken | IF, IHC, WB |
| Cytokeratin 7 | invitrogen | MA1-06315 | Mouse | IF, IHC, WB |
| Cytokeratin 8 | Novusbio | NBP1-48281 | mouse | IF, IHC, WB |
| Cytokeratin 17 | Proteinteach | 18502-1-AP | Rabbit | FC, IF, IHC, IP, WB, ELISA |
| CNN1 | MYBiosource | MBS9435576 | Rabbit | IF, IHC, ICC |
| GAPDH | Thermo Fisher | PA1-9046 | Goat | WB |
| Ki67 | Thermo Fisher | 14-5698-82 | Rat | IF, IHC, WB |
| MUC5AC | Fisher scientific | MS-145-PO | Mouse | IF, IHC, WB |
| MUC5B | Sigma | HPA008246 | Rabbit | IF, IHC |
| MMP1 | Calbiochem | IM35L | Mouse | IF, WB, ELISA |
| MMP2 | ThermoFisher | 436000 | Mouse | IHC, WB, ELISA |
| MMP3 | Calbiochem | IM36L | Mouse | IF, WB, ELISA |
| MMP7 | Bioss antibodies | bs-0423R | Rabbit | WB, ELISA, IHC, IF |
| Muc-1 | Sigma | SAB4200017 | Mouse | IF, WB |
| SFTPC (SPC) | Aviva syst bio | ARP41512_p050 | Rabbit | IF, WB |
| S100A12 | Millipore | AB9728 | Rabbit | IF, WB, ELISA |
| SFTPA | MYBiosource | MBs2028599 | Rabbit | WB, IHC, ICC, IP |
| SFTPD | MYBiosource | MBS2004333 | Rabbit | WB, IHC, ICC, IP |
| SCGB1A1 | Millipore Sigma | ABS1673 | Goat | IF, IHC |
| SCGB3A2 | R&D systems a biotechne brand | AF3545 | Goat | WB, ELISA |
| SCGB3A2 | abcam | ab181853 | Rabbit | WB, ICC/IF |
| Tenascin C | Novusbio | NB110-68136 | mouse | WB, ELISA, IHC |
| TP63 | R&D systems a biotechne | AF1916 | Goat | WB, IF, IHC |
| TP63 | BioCare Medical | CM163A | Mouse | IF, WB |

**Suppl. Table S2. Secondary antibodies used for Immunostaining and Western-blotting assays**

| Protein | Vendor | Product no. | Species of Origin | Species Against | Application & Dilution |
|---|---|---|---|---|---|
| Alexa Fluor 555 donkey anti-mouse lgG(H+L) | Life technologies | A31570 | donkey | Mouse | IF: 1:500 |
| Alexa Fluor 488 donkey anti-mouse IgG (H+L) | Invitrogen | A21202 | Donkey | Mouse | IF: 1:500 |
| Alexa Flour 488 donkey anti-rabbit IgG (H+L) | Invitrogen | A21206 | Donkey | Rabbit | IF: 1:500 |
| Alexa Flour 555 Donkey anti-Rabbit Ig (H+L) | ThermoFisher | A32794 | Donkey | Rabbit | IF: 1:500 |
| Alexa Fluor 647 Donkey Anti-Rabbit IgG (H+L) | Jackson Immuno Research | 711-606-152 | Donkey | Rabbit | IF: 1:500 |
| Alexa Fluor 647 Donkey Anti-Mouse IgG (H+L) | Jackson ImmunoResearch | 715-606-151 | Donkey | Mouse | IF: 1:500 |
| Alexa Fluor 488-Donkey Anti-Chicken | Jackson ImmunoResearch | 703-546-155 | Donkey | Chicken | IF: 1:500 |
| HRP- Donkey Anti-Mouse | Jackson ImmunoResearch | 715-035-151 | Donkey | Mouse | WB: 1:2000 |
| HRP Goat anti-Rabbit | ProteinTech Group | 00001-2 | Goat | Rabbit | WB: 1:2000 |
| HRP Donkey anti-Rabbit | Santa Cruz | sc-2317 | Donkey | Rabbit | WB: 1:2000 |
| Peroxid-conjugated AffiniPure Goat Anti-Chicken | Jackson Immuno Research | 103-035-155 | Goat | Chicken | WB: 1:2000 |
| Donkey anti-Goat (680) | | | Donkey | Goat | WB: 1:10000 |
| Goat anti-Rabbit (800) | | | Goat | Rabbit | WB: 1:10000 |
| Goat anti-Rabbit (680) | | | Goat | Rabbit | WB: 1:10000 |
| Goat anti-Mouse (800) | | | Goat | Mouse | WB: 1:10000 |
| Goat anti-Mouse (680 | | | Goat | Mouse | WB: 1:10000 |
| Donkey Anti-Rabbit IgG | | | Donkey | Rabbit | WB: 1:5000 |
| Donkey Anti-Goat IgG | | | Donkey | Goat | WB: 1:5000 |
| Donkey Anti-Mouse IgG | | | Donkey | Mouse | WB: 1:5000 |
